# A Comparison Of Robust Mendelian Randomization Methods Using Summary Data

**DOI:** 10.1101/577940

**Authors:** Eric A.W. Slob, Stephen Burgess

## Abstract

The number of Mendelian randomization analyses including large numbers of genetic variants is rapidly increasing. This is due to the proliferation of genome-wide association studies, and the desire to obtain more precise estimates of causal effects. Since it is unlikely that all genetic variants will be valid instrumental variables, several robust methods have been proposed. We compare nine robust methods for Mendelian randomization based on summary data that can be implemented using standard statistical software. Methods were compared in three ways: by reviewing their theoretical properties, in an extensive simulation study, and in an empirical example to investigate the effect of body mass index on coronary artery disease risk. In the simulation study, the overall best methods, judged by mean squared error, were the contamination mixture method and the mode based estimation method. These methods generally had well-controlled Type 1 error rates with up to 50% invalid instruments across a range of scenarios. Outlier-robust methods such as MR-Lasso, MR-Robust, and MR-PRESSO, had the narrowest confidence intervals in the empirical example. They performed well when most variants were valid instruments with a few outliers, but less well with several invalid instruments. With isolated exceptions, all methods performed badly when over 50% of the variants were invalid instruments. Our recommendation for investigators is to perform a variety of robust methods that operate in different ways and rely on different assumptions for valid inferences to assess the reliability of Mendelian randomization analyses.

## Introduction

Mendelian randomization (MR) uses genetic variants as instrumental variables (IV) to determine whether an observational association between a risk factor and an outcome is consistent with a causal effect [1, 2]. This approach is less vulnerable to traditional problems of epidemiological studies such as confounding and reverse causality. With the increasing availability of genome-wide association studies that find robust associations between genetic variants and exposures of interest [3, 4], the potential of this approach is rapidly evolving. A genetic variant is a valid IV if (*i*) it is associated with the exposure, (*ii*) it has no direct effect on the outcome, and (*iii*) there are no associations between the variant and any potential confounders.

There has been much discussion on the potentials and limitations of MR, as the IV assumptions cannot be fully tested [1, 5, 6]. Violation of the IV assumptions can lead to invalid conclusions in applied investigations. In practice, the exclusion restriction assumption that the proposed instruments (genetic variants) should not have a direct effect on the outcome of interest is debatable, particularly if the biological roles of the genetic variants are insufficiently understood [5, 7].

Some genetic variants are associated with multiple phenotypic variables [8, 9]. This is referred to as pleiotropy. There are two types of pleiotropy. Vertical pleiotropy occurs when a variant is directly associated with the exposure and another phenotype on the same biological pathway. This does not lead to violation of the IV assumptions provided the only causal pathway from the genetic variant to the outcome passes via the exposure. Horizontal pleiotropy occurs when the second phenotype is on a different biological pathway, and so there may exist different causal pathways from the variant to the outcome. This would violate the exclusion restriction assumption. To solve the problems that arise due to horizontal pleiotropy, several robust methods for MR have been developed that can provide reliable inferences when some genetic variants violate the IV assumptions, or when genetic variants violate the IV assumptions in a particular way. To our knowledge, a comprehensive review and simulation study to compare the statistical performance of these different methods has not been performed.

To focus our simulation study and compare the most relevant robust methods for applied practice, we concentrate on methods that satisfy two criteria. First, the method requires only summary data on estimates (beta-coefficients and standard errors) of genetic variant–exposure and genetic variant– outcome associations. We exclude methods that require individual participant data [10–13], and those that require data on additional variants not associated with the risk factor [14, 15]. This is because the sharing of individual participant data is often impractical, so that many empirical researchers only have access to summary data, and for fairness, to ensure that all methods are using the same information to make inferences. Secondly, the method must be performed using standard statistical software packages. We exclude methods requiring specific computational tools that are unlikely to be accessible to the majority of epidemiologists [16] or are computationally infeasible for large numbers of variants in a reasonable running time [17].

In this article, we review nine robust methods for MR from a theoretical perspective, and evaluate their performance in a simulation study set in a two-sample summary data setting. The methods differ in how they estimate a causal effect of the exposure on the outcome, as well as in the assumptions required for consistent estimation. We consider the weighted median, mode based estimation, MR-PRESSO, MR-Robust, MR-Lasso, MR-Egger, contamination mixture, MR-Mix, and MR-RAPS methods. Some methods take a summarized measure of the variant-specific causal estimates as the overall causal effect estimate (weighted median, and mode based estimation), whereas others remove or downweight outliers (MR-PRESSO, MR-Lasso, MR-Robust), or attempt to model the distribution of the estimates from invalid IVs (MR-Egger, contamination mixture, MR-Mix, and MR-RAPS). We also consider the performance of the methods in an empirical example to evaluate the causal effect of body mass index on coronary artery disease risk.

This paper is organized as follows. First, we give an overview of the robust methods and compare their theoretical properties. Then, we introduce the simulation framework and applied example to compare their properties in practice. Finally, we discuss the implications of this work for applied practice.

## Methods

### Modelling assumptions and summary data

We consider a model as previously described [18, 19] for *J* genetic variants *G*_1_, *G*_2_, *…, G*_*J*_ that are independent in their distributions, a modifiable exposure *X*, an outcome variable *Y*, and a confounder *U*. We assume that all relationships between variables are linear and homogeneous without effect modification, meaning that the same causal effect is estimated by any valid IV [20]. A visual representation of the model is shown in Figure 1.

**Fig. 1:**
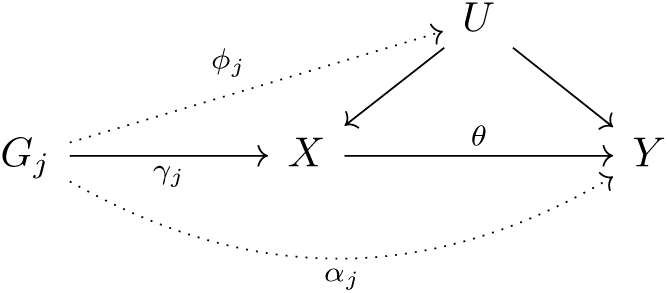
Illustrative diagram showing the model assumed for genetic variant *G*_*j*_, with effect *φ*_*j*_ on the unobserved confounder *U*, effect *γ*_*j*_ on exposure *X*, and direct effect *α*_*j*_ on outcome *Y*. The causal effect of the exposure on the outcome is *θ*. Dotted lines represent possible ways the instrumental variable assumptions could be violated.

We assume that summary data are available on genetic associations with the exposure (betacoefficient 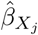 and standard error 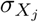) and with the outcome (beta-coefficient 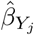 and standard error 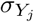) for each variant *G*_*j*_.

### Inverse-variance weighted method

The causal effect of the exposure on the outcome can be estimated using a single genetic variant *G*_*j*_ by the ratio method:

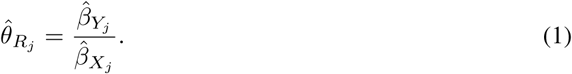

The ratio estimate 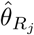 is a consistent estimate of the causal effect if variant *G*_*j*_ satisfies the IV assumptions [20]. If the uncertainty in the genetic association with the exposure is low, then the standard error of the ratio estimate 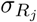 can be approximated as [21]:

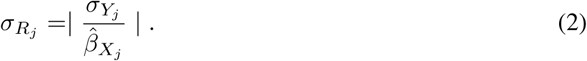

The individual ratio estimates can be combined to obtain a single more efficient estimate. The optimally-efficient combination of the ratio estimates is referred to as the inverse-variance weighted (IVW) estimate [22]:

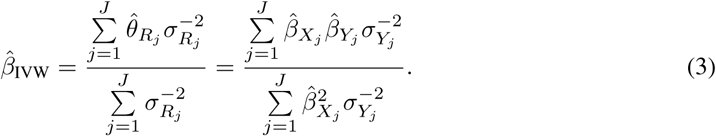

The IVW estimate is equal to the estimate from the two-stage least squares method that is performed using individual participant data [23]. It is a weighted mean of the ratio estimates, where the weights are the inverse-variances of the ratio estimates. The IVW estimate can also be obtained by weighted regression of the genetic associations with the outcome on the genetic associations with the exposure:

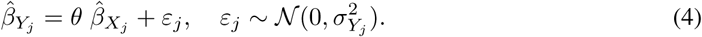

However, the IVW method has a 0% breakdown point, meaning that if only one genetic variant is not a valid IV, then the estimator is typically biased [24]. Bias will be present unless the pleiotropic effects of genetic variants average to zero (balanced pleiotropy) and the pleiotropic effects are independent of the genetic variant–exposure associations (see MR-Egger method below) [19]. With the increasing number of variants used in MR investigations, it is increasingly unlikely that all variants are valid IVs. Hence, it is crucial to consider robust estimation methods despite their lower statistical efficiency (that is, lower power to detect a causal effect).

We proceed to introduce the different robust methods we consider in this study in three categories: consensus methods, outlier-robust methods, and modelling methods.

### Consensus methods

A consensus method is one that takes its causal estimate as a summary measure of the distribution of the ratio estimates. The most straightforward consensus method is the median method. Rather than taking a weighted mean of the ratio estimates as in the IVW method, we take the median of the ratio estimates. The median estimator is consistent (that is, unbiased in large samples) even if up to 50% of the variants are invalid [24]. We consider a weighted version of the median method, where the median is taken from a distribution of the ratio estimates in which genetic variants with more precise ratio estimates receive more weight. Here, an unbiased estimate will be obtained if up to 50% of the weight comes from variants that are valid IVs. We refer to this as the ‘majority valid’ assumption.

A related assumption is the ‘plurality valid’ assumption [11]. In large samples, while ratio estimates for all valid IVs should equal the true causal effect, ratio estimates for invalid IVs will take different values. The ‘plurality valid’ assumption is that, out of all the different values taken by ratio estimates in large samples (we term these the ratio estimands), the true causal effect is the value taken for the largest number of genetic variants (that is, the modal ratio estimand). For example, the plurality assumption would be satisfied if only 40% of the genetic variants are valid instruments, provided that out of the remaining 60% invalid instruments, no larger group with the same ratio estimand exists. This assumption is also referred to as the Zero Modal Pleiotropy Assumption (ZEMPA) [25].

This assumption is exploited by the mode based estimation (MBE) method [25]. As no two ratio estimates will be identical in finite samples, it is not possible to take the mode of the ratio estimates directly. In the MBE method, a normal density is drawn for each genetic variant centered at its ratio estimate. The spread of this density depends on a bandwidth parameter, and (for the weighted version of the MBE method) the precision of the ratio estimate. A smoothed density function is then constructed by summing these normal densities. The maximum of this distribution is the causal estimate.

As these consensus methods take the median or mode of the ratio estimate distribution as the causal estimate, they are naturally robust to outliers, as the median and mode of a distribution are unaffected by the magnitude of extreme values. However, they are still influenced by outliers, as these variants still contribute to determining the location of the median or mode of a distribution. These methods can also be sensitive to changes in the ratio estimates for variants that contribute to the median or mode, and to the addition and removal of variants from the analysis. Additionally, the methods may not be as efficient as those that base their estimates on all the genetic variants.

### Outlier-robust methods

Next, we present three outlier-robust methods. These methods either downweight or remove genetic variants from the analysis that have outlying ratio estimates. They provide consistent estimates under the same assumptions as the IVW method for the set of genetic variants that are not identified as outliers.

In the MR-Pleiotropy Residual Sum and Outlier (MR-PRESSO) method [26], the IVW method is implemented by regression using all the genetic variants, and the residual sum of squares (RSS) is calculated from the regression equation. The RSS is a heterogeneity measure for the ratio estimates. Then, the IVW method is performed omitting each genetic variant from the analysis in turn. If the RSS decreases substantially compared to a simulated expected distribution, then that variant is removed from the analysis. This procedure is repeated until no further variants are removed from the analysis. The causal estimate is then obtained by the IVW method using the remaining genetic variants.

In MR-Robust, the IVW method is performed by regression, except that instead of using ordinary least squares regression, MM-estimation is used combined with Tukey’s biweight loss function [27]. MM-estimation provides robustness against influential points and Tukey’s loss function provides robustness against outliers. Tukey’s loss function is a truncated quadratic function, meaning that there is a limit in the degree to which an outlier contributes to the analysis [28]. This contrasts with the quadratic loss function used in ordinary least squares regression, which is unbounded, meaning that a single outlier can have an unlimited effect on the IVW estimate.

In MR-Lasso, the IVW regression model is augmented by adding an intercept term for each genetic variant [27]. The IVW estimate is the value of *θ* that minimizes:

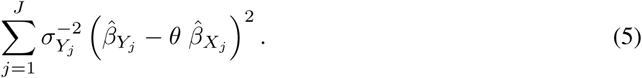

In MR-Lasso, we minimize:

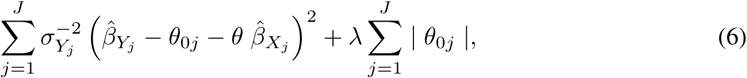

where *λ* is a tuning parameter. As the regression equation contains more parameters than there are genetic variants, a lasso penalty term is added for identification [29]. The intercept term 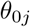 represents the direct (pleiotropic) effect on the outcome, and should be zero for a valid IV, but will be non-zero for an invalid IV. The causal estimate is then obtained by the IVW method using the genetic variants that had 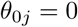 in equation (6). A heterogeneity criterion is used to determine the value of *λ*. Increasing *λ* means that more of the pleiotropy parameters equal zero and so the corresponding variants are included in the analysis; we increase *λ* step-by-step until one step before there is more heterogeneity in the ratio estimates for variants included in the analysis than expected by chance alone.

The MR-PRESSO and MR-Lasso methods remove variants from the analysis, whereas MR-Robust downweights variants. These methods will be valuable when there is a small number of genetic variants with heterogeneous ratio estimates, as they will be removed from the analysis or heavily down-weighted, and so will not influence the overall estimate. In such a case, these methods are likely to be efficient, as they are based on the IVW method. The methods are less likely to be valuable when there is a larger number of genetic variants that are pleiotropic, particularly if the pleiotropic effects are small in magnitude, and when the average pleiotropic effect of non-outliers is not zero.

### Modelling methods

Finally, we present four methods that attempt to model the distribution of estimates from invalid IVs or make a specific assumption about the way in which the IV assumptions are violated. The MR-Egger method is performed similarly to the IVW method, except that the regression model contains an intercept term *θ*_0_:

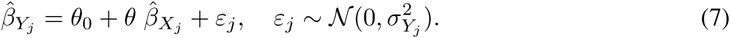

This differs from the MR-Lasso method, as there is only one intercept term, which represents the average pleiotropic effect. The MR-Egger method gives consistent estimates of the causal effect under the Instrument Strength Independent of Direct Effect (InSIDE) assumption, which states that pleiotropic effects of genetic variants must be uncorrelated with genetic variant–exposure association. As the regression model is no longer symmetric to changes in the signs of the genetic association estimates (which result from switching the reference and effect alleles), we first re-orientate the genetic associations before performing the regression by fixing all genetic associations with the exposure to be positive, and correspondingly changing the signs of the genetic associations with the outcome if necessary. The intercept in MR-Egger also provides a test of the IV assumptions. The intercept will differ from zero when either the average pleiotropic effect is not zero, or the InSIDE assumption is violated. These are precisely the conditions required for the IVW estimate to be unbiased.

The contamination mixture method assumes that only some of the genetic variants are valid IVs [30]. We construct a likelihood function from the ratio estimates. If a variant is a valid instrument, then its ratio estimate is assumed to be normally distributed about the true causal effect *θ* with variance 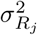. If a variant is not a valid instrument, then its ratio estimate is assumed to be normally distributed about zero with variance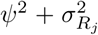, where *ψ*^2^ represents the variance of the estimands from invalid IVs. This parameter is specified by the analyst. We then maximize the likelihood over different values of the causal effect *θ* and different configurations of valid and invalid IVs. Maximization is performed in linear time by first constructing a profile likelihood as a function of *θ*, and then maximizing this function with respect to *θ*. The value of *θ* that maximizes the profile likelihood is the causal estimate.

The MR-Mix method [31] is similar to the contamination mixture method, except that rather than dividing the genetic variants into valid and invalid IVs, the method divides variants into four categories: 1) variants that directly influence the exposure only (valid instruments), and 2) variants that influence the exposure and outcome, 3) that influence the outcome only, and 4) that neither influence the exposure or outcome (invalid instruments). This allows for more flexibility in modelling genetic variants, although potentially leads to more uncertainty in assigning genetic variants to categories.

The MR-Robust Adjusted Profile Score (RAPS) [32] method models the pleiotropic effects of genetic variants directly using a random-effects distribution. The pleiotropic effects are assumed to be normally distributed about zero with unknown variance. Estimates are obtained using a profile likelihood function for the causal effect and the variance of the pleiotropic effect distribution. To provide further robustness to outliers, either Tukey’s biweight loss function or Huber’s loss function [28] can be used.

Modelling methods are likely to be valuable when the modelling assumptions are correct, but not when the assumptions are incorrect. For example, the MR-Egger method requires the InSIDE assumption to be satisfied to give a consistent estimate. The MR-RAPS method is likely to perform well when pleiotropic effects truly are normally distributed about zero, but less well when they are not. The MR-Mix method is likely to require large numbers of genetic variants in order to correct classify variants into the different categories. The contamination mixture method is less likely to be affected by modelling assumptions as it does not make such strict assumptions, but it is likely to be sensitive to specification of the variance parameter.

### Simulation study

To compare the performance of these methods in a realistic setting, we perform a simulation study. Full details of the simulation study are given in the Supplementary Material. In brief, we consider three scenarios:

1. balanced pleiotropy, InSIDE satisfied – invalid IVs have direct effects on the outcome generated from a normal distribution centered at zero;
2. directional pleiotropy, InSIDE satisfied – invalid IVs have direct effects on the outcome generated from a normal distribution centered away from zero;
3. directional pleiotropy, InSIDE violated – invalid IVs have direct effects on the outcome generated from a normal distribution centered away from zero, and indirect effects on the outcome via the confounder.

We simulated data on *J* = 10, 30, and 100 genetic variants. A portion of the genetic variants were invalid IVs (30%, 50% and 70%), and the direct effects of the variants explain 10% of the variance in the exposure. Summary genetic associations were calculated for the exposure and the outcome on non-overlapping sets of individuals, each consisting of 10 000 individuals [33]. This situation is often referred to as two-sample summary data MR [34]. We considered situations with a null causal effect (*θ* = 0) and a positive causal effect (*θ* = 0.2). In total, 10 000 datasets were generated in each scenario. We additionally considered scenarios with 500 genetic variants and a wider range of proportions of invalid IVs (additionally 1%, 5%, and 10% invalid).

### Empirical example: the effect of body mass index on coronary artery disease risk

We also compare the methods in an empirical example considering the effect of body mass index (BMI) on coronary artery disease (CAD) risk. Since BMI is influenced by several biological mechanisms [35], it is likely that the exclusion restriction is not satisfied for all associated genetic variants. Hence it is necessary to use robust methods to analyse these data. Additionally, we consider methods that detect outliers (MR-Presso, MR-Robust, MR-Lasso, contamination mixture, MR-Mix, and MR-RAPS), and compare whether the same outliers are detected in each of these methods.

We take 97 genome-wide significant variants associated with BMI from the GIANT consortium [36]. Associations with BMI are estimated in up to 339,224 participants from this consortium. Associations with coronary artery disease risk are estimated in up to 60,801 CAD cases and 123,504 controls from the CARDIoGRAMplusC4D Consortium [37]. Association estimates for CAD were available for 94 of these variants.

The scatter plot of the genetic associations with BMI and CAD risk is shown in Figure 2. While most variants seem to suggest a harmful effect of increased BMI on CAD risk, there is substantial heterogeneity in the plot. This suggests that some of the variants violate the IV assumptions.

**Fig. 2:**
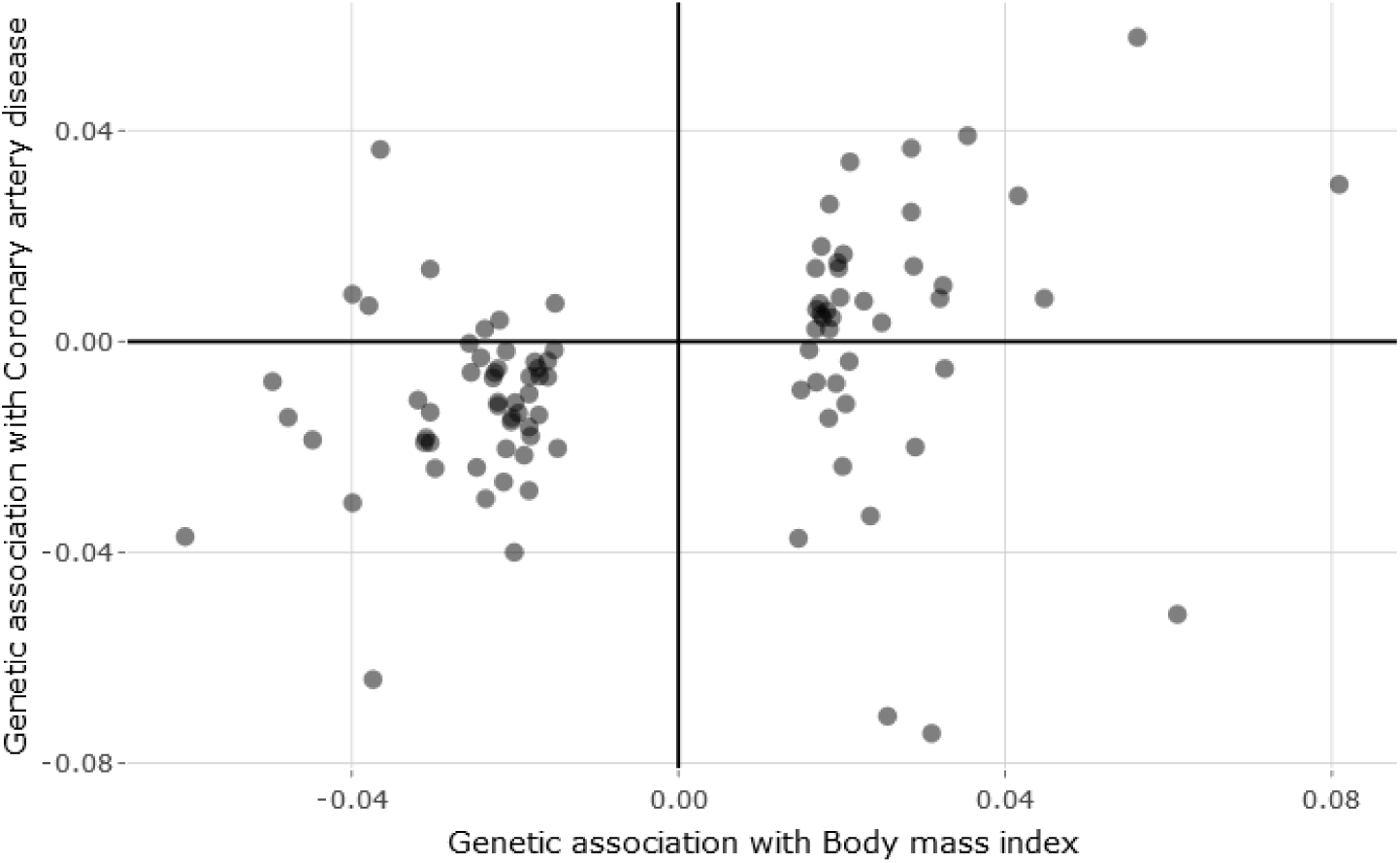
Scatter plot of genetic associations with BMI (standard deviation units) and coronary artery disease risk (log odds ratios) for 94 variants taken from the GIANT and CARDIoGRAMplusC4D consortia respectively.

## Results

### Simulation study

Results of the simulation study are presented in Table 1 (10 variants), Table 2 (30 variants), and Table 3 (100 variants). For each scenario, we present the mean, median, and standard deviation of estimates across simulations, and the empirical power of the 95% confidence interval. With a null causal effect, the empirical power is the Type 1 error rate, and should be close to 0.05. The mean squared error across simulations for the different methods with a null causal effect is presented in Figure 3 (Scenario 2), and Figure 4 (Scenario 3) for 30 variants. The corresponding plots for 10 variants (Supplementary Figures 5 and 6) and 100 variants (Supplementary Figures 7 and 8) were broadly similar, as were results with 500 variants (Supplementary Tables 6 and 7).

**Table 1:**
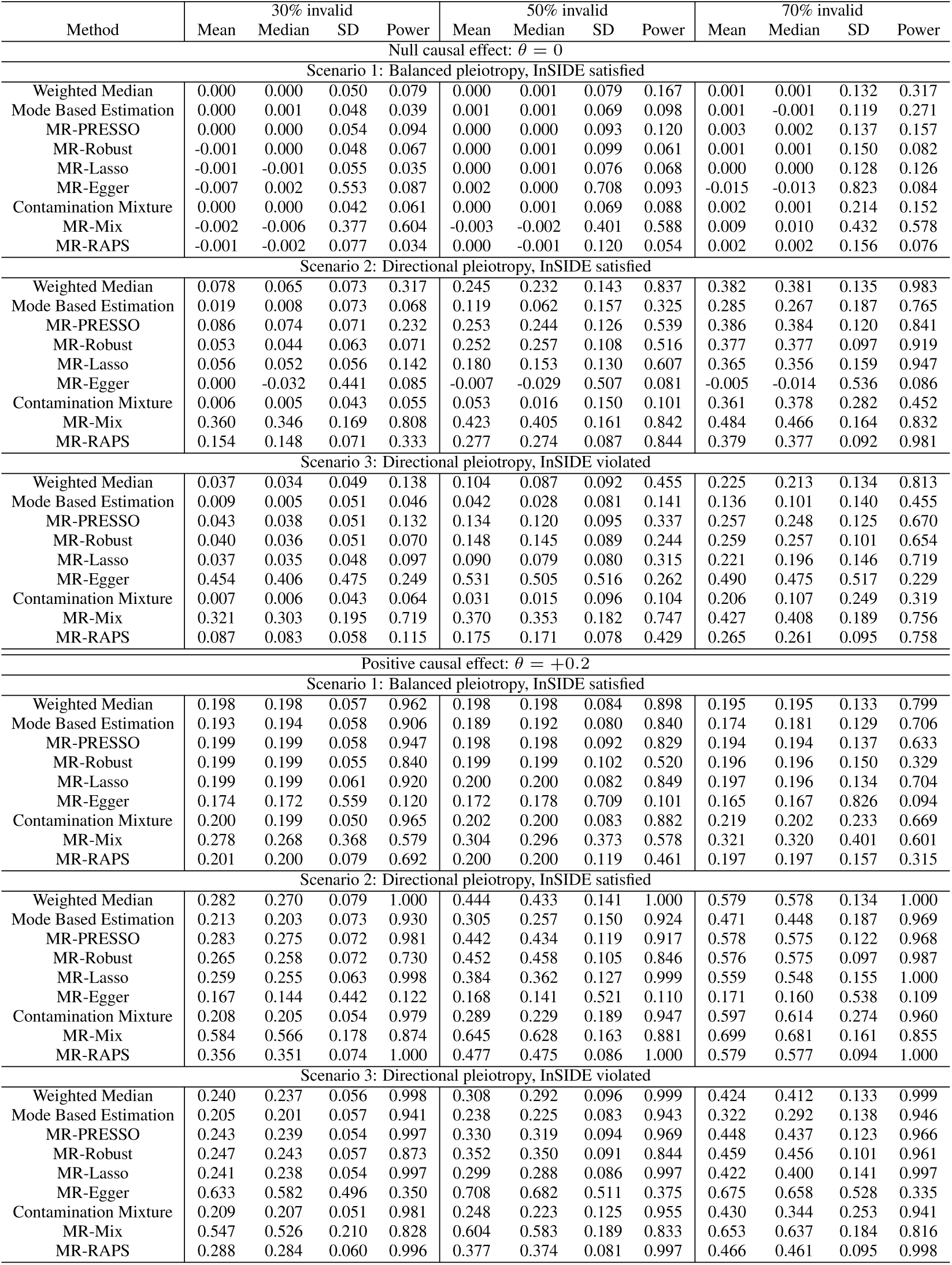
Mean, median, standard deviation (SD) of estimates, and empirical power with 10 genetic variants.

**Table 2:** Mean, median, standard deviation (SD) of estimates, and empirical power with 30 genetic variants.

| Method | 30% invalid |  |  |  | 50% invalid |  |  |  | 70% invalid |  |  |  |
| --- | --- | --- | --- | --- | --- | --- | --- | --- | --- | --- | --- | --- |
|  | Mean | Median | SD | Power | Mean | Median | SD | Power | Mean | Median | SD | Power |
| Null causal effect: $\theta = 0$ | | | | | | | | | | | | |
| Scenario 1: Balanced pleiotropy, InSIDE satisfied |  |  |  |  |  |  |  |  |  |  |  |  |
| Weighted Median | 0.000 | 0.000 | 0.032 | 0.058 | 0.000 | 0.000 | 0.044 | 0.112 | 0.000 | 0.000 | 0.068 | 0.208 |
| Mode Based Estimation | 0.000 | 0.000 | 0.035 | 0.014 | 0.000 | 0.001 | 0.038 | 0.033 | 0.000 | 0.000 | 0.058 | 0.111 |
| MR-PRESSO | 0.000 | 0.000 | 0.029 | 0.090 | 0.001 | 0.001 | 0.046 | 0.152 | 0.000 | 0.000 | 0.072 | 0.233 |
| MR-Robust | 0.000 | 0.000 | 0.030 | 0.047 | 0.001 | 0.001 | 0.059 | 0.034 | -0.001 | -0.002 | 0.098 | 0.052 |
| MR-Lasso | 0.000 | 0.000 | 0.029 | 0.045 | 0.000 | 0.001 | 0.041 | 0.072 | 0.000 | 0.000 | 0.067 | 0.123 |
| MR-Egger | 0.004 | 0.004 | 0.451 | 0.067 | 0.005 | 0.010 | 0.563 | 0.060 | 0.008 | 0.011 | 0.673 | 0.063 |
| Contamination Mixture | 0.000 | 0.000 | 0.028 | 0.067 | 0.000 | 0.001 | 0.038 | 0.094 | 0.000 | 0.000 | 0.071 | 0.153 |
| MR-Mix | -0.004 | -0.002 | 0.206 | 0.202 | 0.002 | 0.000 | 0.307 | 0.235 | 0.002 | 0.000 | 0.411 | 0.302 |
| MR-RAPS | 0.000 | 0.000 | 0.047 | 0.018 | 0.001 | 0.000 | 0.077 | 0.041 | -0.001 | -0.001 | 0.104 | 0.052 |
| Scenario 2: Directional pleiotropy, InSIDE satisfied |  |  |  |  |  |  |  |  |  |  |  |  |
| Weighted Median | 0.085 | 0.082 | 0.042 | 0.653 | 0.293 | 0.289 | 0.106 | 0.994 | 0.452 | 0.452 | 0.089 | 1.000 |
| Mode Based Estimation | 0.006 | 0.005 | 0.034 | 0.019 | 0.053 | 0.037 | 0.082 | 0.120 | 0.302 | 0.285 | 0.189 | 0.742 |
| MR-PRESSO | 0.093 | 0.090 | 0.044 | 0.749 | 0.278 | 0.273 | 0.077 | 0.999 | 0.438 | 0.437 | 0.078 | 1.000 |
| MR-Robust | 0.049 | 0.045 | 0.041 | 0.044 | 0.289 | 0.294 | 0.075 | 0.691 | 0.433 | 0.433 | 0.059 | 0.999 |
| MR-Lasso | 0.037 | 0.036 | 0.030 | 0.292 | 0.183 | 0.171 | 0.084 | 0.963 | 0.419 | 0.417 | 0.106 | 1.000 |
| MR-Egger | 0.000 | -0.021 | 0.343 | 0.058 | 0.000 | -0.013 | 0.405 | 0.062 | 0.004 | -0.010 | 0.432 | 0.065 |
| Contamination Mixture | 0.005 | 0.005 | 0.028 | 0.070 | 0.017 | 0.013 | 0.051 | 0.106 | 0.373 | 0.432 | 0.290 | 0.592 |
| MR-Mix | 0.254 | 0.236 | 0.192 | 0.455 | 0.384 | 0.367 | 0.208 | 0.548 | 0.513 | 0.511 | 0.190 | 0.634 |
| MR-RAPS | 0.175 | 0.174 | 0.047 | 0.950 | 0.321 | 0.320 | 0.056 | 1.000 | 0.435 | 0.434 | 0.058 | 1.000 |
| Scenario 3: Directional pleiotropy, InSIDE violated |  |  |  |  |  |  |  |  |  |  |  |  |
| Weighted Median | 0.042 | 0.041 | 0.031 | 0.243 | 0.106 | 0.101 | 0.055 | 0.726 | 0.256 | 0.250 | 0.096 | 0.980 |
| Mode Based Estimation | 0.004 | 0.003 | 0.034 | 0.014 | 0.021 | 0.018 | 0.039 | 0.048 | 0.089 | 0.072 | 0.088 | 0.331 |
| MR-PRESSO | 0.045 | 0.044 | 0.032 | 0.335 | 0.140 | 0.136 | 0.056 | 0.893 | 0.281 | 0.277 | 0.082 | 0.997 |
| MR-Robust | 0.039 | 0.038 | 0.033 | 0.071 | 0.163 | 0.163 | 0.063 | 0.409 | 0.301 | 0.301 | 0.065 | 0.968 |
| MR-Lasso | 0.025 | 0.024 | 0.028 | 0.178 | 0.082 | 0.078 | 0.048 | 0.658 | 0.238 | 0.228 | 0.100 | 0.980 |
| MR-Egger | 0.838 | 0.817 | 0.375 | 0.754 | 0.919 | 0.904 | 0.365 | 0.789 | 0.845 | 0.830 | 0.359 | 0.719 |
| Contamination Mixture | 0.006 | 0.006 | 0.028 | 0.072 | 0.017 | 0.015 | 0.040 | 0.123 | 0.139 | 0.061 | 0.203 | 0.344 |
| MR-Mix | 0.205 | 0.184 | 0.166 | 0.360 | 0.299 | 0.283 | 0.178 | 0.436 | 0.409 | 0.395 | 0.191 | 0.516 |
| MR-RAPS | 0.097 | 0.096 | 0.037 | 0.499 | 0.202 | 0.201 | 0.052 | 0.963 | 0.307 | 0.306 | 0.061 | 0.999 |
| Positive causal effect: $\theta = +0.2$ | | | | | | | | | | | | |
| Scenario 1: Balanced pleiotropy, InSIDE satisfied |  |  |  |  |  |  |  |  |  |  |  |  |
| Weighted Median | 0.196 | 0.195 | 0.036 | 1.000 | 0.195 | 0.194 | 0.048 | 0.991 | 0.197 | 0.196 | 0.073 | 0.942 |
| Mode Based Estimation | 0.189 | 0.189 | 0.041 | 0.976 | 0.187 | 0.187 | 0.044 | 0.964 | 0.180 | 0.181 | 0.067 | 0.853 |
| MR-PRESSO | 0.198 | 0.197 | 0.033 | 1.000 | 0.197 | 0.197 | 0.049 | 0.989 | 0.199 | 0.197 | 0.075 | 0.929 |
| MR-Robust | 0.198 | 0.197 | 0.035 | 0.995 | 0.197 | 0.197 | 0.061 | 0.824 | 0.199 | 0.198 | 0.098 | 0.518 |
| MR-Lasso | 0.198 | 0.198 | 0.033 | 1.000 | 0.197 | 0.197 | 0.045 | 0.996 | 0.199 | 0.199 | 0.073 | 0.939 |
| MR-Egger | 0.113 | 0.120 | 0.453 | 0.076 | 0.118 | 0.123 | 0.566 | 0.073 | 0.116 | 0.121 | 0.666 | 0.064 |
| Contamination Mixture | 0.199 | 0.199 | 0.033 | 1.000 | 0.199 | 0.198 | 0.043 | 0.993 | 0.206 | 0.201 | 0.084 | 0.933 |
| MR-Mix | 0.255 | 0.225 | 0.213 | 0.378 | 0.276 | 0.232 | 0.283 | 0.367 | 0.323 | 0.254 | 0.385 | 0.386 |
| MR-RAPS | 0.200 | 0.200 | 0.049 | 0.942 | 0.200 | 0.200 | 0.077 | 0.708 | 0.201 | 0.200 | 0.103 | 0.502 |
| Scenario 2: Directional pleiotropy, InSIDE satisfied |  |  |  |  |  |  |  |  |  |  |  |  |
| Weighted Median | 0.293 | 0.290 | 0.048 | 1.000 | 0.493 | 0.488 | 0.104 | 1.000 | 0.646 | 0.647 | 0.091 | 1.000 |
| Mode Based Estimation | 0.198 | 0.196 | 0.040 | 0.987 | 0.246 | 0.232 | 0.081 | 0.940 | 0.472 | 0.434 | 0.188 | 0.938 |
| MR-PRESSO | 0.294 | 0.292 | 0.045 | 1.000 | 0.465 | 0.461 | 0.072 | 1.000 | 0.624 | 0.622 | 0.080 | 1.000 |
| MR-Robust | 0.265 | 0.262 | 0.049 | 0.941 | 0.495 | 0.500 | 0.073 | 0.958 | 0.632 | 0.632 | 0.061 | 1.000 |
| MR-Lasso | 0.243 | 0.242 | 0.036 | 1.000 | 0.396 | 0.386 | 0.084 | 1.000 | 0.614 | 0.610 | 0.104 | 1.000 |
| MR-Egger | 0.110 | 0.096 | 0.352 | 0.081 | 0.110 | 0.100 | 0.401 | 0.076 | 0.111 | 0.103 | 0.429 | 0.077 |
| Contamination Mixture | 0.206 | 0.204 | 0.034 | 1.000 | 0.233 | 0.218 | 0.093 | 0.999 | 0.648 | 0.707 | 0.270 | 0.999 |
| MR-Mix | 0.504 | 0.490 | 0.171 | 0.647 | 0.614 | 0.601 | 0.173 | 0.672 | 0.718 | 0.708 | 0.166 | 0.708 |
| MR-RAPS | 0.378 | 0.377 | 0.048 | 1.000 | 0.522 | 0.522 | 0.057 | 1.000 | 0.636 | 0.636 | 0.060 | 1.000 |
| Scenario 3: Directional pleiotropy, InSIDE violated |  |  |  |  |  |  |  |  |  |  |  |  |
| Weighted Median | 0.242 | 0.241 | 0.036 | 1.000 | 0.312 | 0.307 | 0.058 | 1.000 | 0.453 | 0.448 | 0.095 | 1.000 |
| Mode Based Estimation | 0.195 | 0.193 | 0.039 | 0.994 | 0.213 | 0.211 | 0.043 | 0.995 | 0.277 | 0.263 | 0.086 | 0.966 |
| MR-PRESSO | 0.247 | 0.246 | 0.035 | 1.000 | 0.341 | 0.339 | 0.056 | 1.000 | 0.473 | 0.469 | 0.079 | 1.000 |
| MR-Robust | 0.246 | 0.245 | 0.038 | 0.997 | 0.370 | 0.370 | 0.062 | 0.982 | 0.500 | 0.500 | 0.066 | 0.999 |
| MR-Lasso | 0.228 | 0.227 | 0.032 | 1.000 | 0.294 | 0.290 | 0.054 | 1.000 | 0.441 | 0.433 | 0.095 | 1.000 |
| MR-Egger | 0.987 | 0.964 | 0.375 | 0.859 | 1.081 | 1.063 | 0.368 | 0.890 | 0.993 | 0.975 | 0.354 | 0.832 |
| Contamination Mixture | 0.207 | 0.206 | 0.034 | 1.000 | 0.225 | 0.221 | 0.054 | 0.999 | 0.399 | 0.302 | 0.233 | 0.997 |
| MR-Mix | 0.455 | 0.434 | 0.169 | 0.592 | 0.549 | 0.535 | 0.168 | 0.602 | 0.637 | 0.623 | 0.171 | 0.637 |
| MR-RAPS | 0.298 | 0.297 | 0.039 | 1.000 | 0.403 | 0.402 | 0.053 | 1.000 | 0.508 | 0.507 | 0.063 | 1.000 |

**Table 3:** Mean, median, standard deviation (SD) of estimates, and empirical power with 100 genetic variants.

| Method | 30% invalid |  |  |  | 50% invalid |  |  |  | 70% invalid |  |  |  |
| --- | --- | --- | --- | --- | --- | --- | --- | --- | --- | --- | --- | --- |
|  | Mean | Median | SD | Power | Mean | Median | SD | Power | Mean | Median | SD | Power |
| Null causal effect: $\theta = 0$ | | | | | | | | | | | | |
| Scenario 1: Balanced pleiotropy, InSIDE satisfied |  |  |  |  |  |  |  |  |  |  |  |  |
| Weighted Median | 0.000 | 0.000 | 0.019 | 0.057 | 0.000 | 0.000 | 0.026 | 0.098 | 0.000 | 0.000 | 0.038 | 0.176 |
| Mode Based Estimation | 0.000 | 0.000 | 0.026 | 0.005 | 0.000 | 0.000 | 0.026 | 0.010 | 0.000 | 0.000 | 0.033 | 0.036 |
| MR-PRESSO | 0.000 | 0.000 | 0.017 | 0.092 | 0.000 | 0.000 | 0.027 | 0.151 | 0.000 | 0.000 | 0.041 | 0.229 |
| MR-Robust | 0.000 | 0.000 | 0.017 | 0.044 | 0.000 | 0.000 | 0.033 | 0.035 | 0.000 | 0.000 | 0.055 | 0.044 |
| MR-Lasso | 0.000 | 0.000 | 0.016 | 0.047 | 0.000 | 0.000 | 0.022 | 0.080 | 0.000 | 0.000 | 0.034 | 0.130 |
| MR-Egger | 0.000 | 0.000 | 0.257 | 0.054 | 0.000 | 0.003 | 0.327 | 0.055 | -0.002 | -0.001 | 0.385 | 0.055 |
| Contamination Mixture | 0.000 | 0.000 | 0.017 | 0.070 | 0.000 | 0.000 | 0.023 | 0.100 | 0.000 | 0.001 | 0.037 | 0.164 |
| MR-Mix | 0.000 | 0.000 | 0.062 | 0.042 | 0.000 | 0.000 | 0.068 | 0.035 | 0.000 | 0.000 | 0.085 | 0.037 |
| MR-RAPS | 0.000 | 0.000 | 0.027 | 0.018 | 0.000 | 0.000 | 0.044 | 0.035 | 0.000 | 0.000 | 0.059 | 0.046 |
| Scenario 2: Directional pleiotropy, InSIDE satisfied |  |  |  |  |  |  |  |  |  |  |  |  |
| Weighted Median | 0.092 | 0.091 | 0.024 | 0.990 | 0.306 | 0.305 | 0.063 | 1.000 | 0.471 | 0.471 | 0.050 | 1.000 |
| Mode Based Estimation | 0.005 | 0.004 | 0.024 | 0.008 | 0.028 | 0.027 | 0.026 | 0.043 | 0.246 | 0.178 | 0.177 | 0.590 |
| MR-PRESSO | 0.095 | 0.094 | 0.024 | 0.995 | 0.269 | 0.268 | 0.040 | 1.000 | 0.437 | 0.436 | 0.044 | 1.000 |
| MR-Robust | 0.053 | 0.052 | 0.024 | 0.178 | 0.307 | 0.308 | 0.041 | 0.990 | 0.450 | 0.450 | 0.034 | 1.000 |
| MR-Lasso | 0.042 | 0.041 | 0.019 | 0.767 | 0.206 | 0.201 | 0.051 | 1.000 | 0.446 | 0.447 | 0.056 | 1.000 |
| MR-Egger | -0.002 | -0.002 | 0.195 | 0.049 | -0.003 | -0.005 | 0.228 | 0.052 | 0.002 | 0.003 | 0.244 | 0.054 |
| Contamination Mixture | 0.006 | 0.006 | 0.017 | 0.087 | 0.017 | 0.016 | 0.025 | 0.179 | 0.391 | 0.492 | 0.277 | 0.793 |
| MR-Mix | 0.083 | 0.030 | 0.163 | 0.134 | 0.235 | 0.098 | 0.272 | 0.273 | 0.468 | 0.532 | 0.282 | 0.452 |
| MR-RAPS | 0.183 | 0.182 | 0.027 | 1.000 | 0.335 | 0.335 | 0.032 | 1.000 | 0.454 | 0.453 | 0.034 | 1.000 |
| Scenario 3: Directional pleiotropy, InSIDE violated |  |  |  |  |  |  |  |  |  |  |  |  |
| Weighted Median | 0.044 | 0.044 | 0.018 | 0.651 | 0.109 | 0.108 | 0.029 | 0.993 | 0.265 | 0.264 | 0.056 | 1.000 |
| Mode Based Estimation | 0.003 | 0.003 | 0.024 | 0.006 | 0.014 | 0.013 | 0.023 | 0.022 | 0.056 | 0.053 | 0.037 | 0.242 |
| MR-PRESSO | 0.048 | 0.048 | 0.018 | 0.801 | 0.140 | 0.139 | 0.030 | 1.000 | 0.280 | 0.279 | 0.044 | 1.000 |
| MR-Robust | 0.040 | 0.040 | 0.019 | 0.313 | 0.170 | 0.169 | 0.036 | 0.945 | 0.315 | 0.315 | 0.037 | 1.000 |
| MR-Lasso | 0.026 | 0.026 | 0.017 | 0.485 | 0.090 | 0.088 | 0.028 | 0.991 | 0.258 | 0.255 | 0.059 | 1.000 |
| MR-Egger | 0.902 | 0.902 | 0.193 | 0.998 | 0.980 | 0.980 | 0.185 | 1.000 | 0.882 | 0.881 | 0.181 | 0.998 |
| Contamination Mixture | 0.007 | 0.006 | 0.016 | 0.096 | 0.018 | 0.017 | 0.023 | 0.202 | 0.071 | 0.049 | 0.105 | 0.481 |
| MR-Mix | 0.064 | 0.035 | 0.119 | 0.122 | 0.163 | 0.080 | 0.207 | 0.202 | 0.325 | 0.315 | 0.262 | 0.323 |
| MR-RAPS | 0.100 | 0.099 | 0.021 | 0.993 | 0.211 | 0.210 | 0.030 | 1.000 | 0.322 | 0.322 | 0.035 | 1.000 |
| Positive causal effect: $\theta = +0.2$ | | | | | | | | | | | | |
| Scenario 1: Balanced pleiotropy, InSIDE satisfied |  |  |  |  |  |  |  |  |  |  |  |  |
| Weighted Median | 0.192 | 0.192 | 0.022 | 1.000 | 0.193 | 0.192 | 0.029 | 1.000 | 0.193 | 0.193 | 0.041 | 0.999 |
| Mode Based Estimation | 0.180 | 0.180 | 0.031 | 0.999 | 0.180 | 0.180 | 0.030 | 0.999 | 0.175 | 0.175 | 0.038 | 0.986 |
| MR-PRESSO | 0.196 | 0.195 | 0.020 | 1.000 | 0.197 | 0.196 | 0.030 | 1.000 | 0.197 | 0.196 | 0.045 | 0.999 |
| MR-Robust | 0.196 | 0.196 | 0.020 | 1.000 | 0.196 | 0.196 | 0.035 | 0.998 | 0.197 | 0.197 | 0.057 | 0.925 |
| MR-Lasso | 0.196 | 0.196 | 0.019 | 1.000 | 0.196 | 0.197 | 0.025 | 1.000 | 0.196 | 0.196 | 0.038 | 1.000 |
| MR-Egger | 0.047 | 0.048 | 0.256 | 0.062 | 0.051 | 0.054 | 0.329 | 0.062 | 0.045 | 0.049 | 0.386 | 0.055 |
| Contamination Mixture | 0.196 | 0.196 | 0.020 | 1.000 | 0.198 | 0.198 | 0.027 | 1.000 | 0.199 | 0.198 | 0.043 | 0.999 |
| MR-Mix | 0.210 | 0.208 | 0.076 | 0.293 | 0.211 | 0.200 | 0.095 | 0.401 | 0.215 | 0.200 | 0.134 | 0.626 |
| MR-RAPS | 0.200 | 0.200 | 0.028 | 1.000 | 0.200 | 0.199 | 0.044 | 0.989 | 0.201 | 0.201 | 0.060 | 0.910 |
| Scenario 2: Directional pleiotropy, InSIDE satisfied |  |  |  |  |  |  |  |  |  |  |  |  |
| Weighted Median | 0.301 | 0.300 | 0.029 | 1.000 | 0.505 | 0.503 | 0.060 | 1.000 | 0.662 | 0.662 | 0.052 | 1.000 |
| Mode Based Estimation | 0.189 | 0.187 | 0.028 | 1.000 | 0.217 | 0.215 | 0.034 | 0.984 | 0.425 | 0.370 | 0.168 | 0.892 |
| MR-PRESSO | 0.305 | 0.305 | 0.027 | 1.000 | 0.471 | 0.471 | 0.040 | 1.000 | 0.627 | 0.626 | 0.043 | 1.000 |
| MR-Robust | 0.274 | 0.273 | 0.031 | 1.000 | 0.512 | 0.513 | 0.039 | 1.000 | 0.646 | 0.646 | 0.035 | 1.000 |
| MR-Lasso | 0.251 | 0.251 | 0.023 | 1.000 | 0.421 | 0.418 | 0.051 | 1.000 | 0.636 | 0.636 | 0.056 | 1.000 |
| MR-Egger | 0.053 | 0.050 | 0.201 | 0.068 | 0.053 | 0.052 | 0.231 | 0.061 | 0.046 | 0.047 | 0.247 | 0.058 |
| Contamination Mixture | 0.206 | 0.205 | 0.022 | 1.000 | 0.226 | 0.223 | 0.037 | 1.000 | 0.702 | 0.761 | 0.217 | 1.000 |
| MR-Mix | 0.354 | 0.302 | 0.184 | 0.460 | 0.540 | 0.525 | 0.251 | 0.544 | 0.730 | 0.753 | 0.220 | 0.605 |
| MR-RAPS | 0.386 | 0.386 | 0.029 | 1.000 | 0.536 | 0.536 | 0.033 | 1.000 | 0.654 | 0.654 | 0.036 | 1.000 |
| Scenario 3: Directional pleiotropy, InSIDE violated |  |  |  |  |  |  |  |  |  |  |  |  |
| Weighted Median | 0.244 | 0.243 | 0.022 | 1.000 | 0.316 | 0.315 | 0.034 | 1.000 | 0.461 | 0.459 | 0.055 | 1.000 |
| Mode Based Estimation | 0.187 | 0.186 | 0.028 | 1.000 | 0.201 | 0.199 | 0.028 | 1.000 | 0.244 | 0.240 | 0.044 | 0.997 |
| MR-PRESSO | 0.255 | 0.255 | 0.022 | 1.000 | 0.349 | 0.348 | 0.032 | 1.000 | 0.478 | 0.477 | 0.043 | 1.000 |
| MR-Robust | 0.249 | 0.249 | 0.023 | 1.000 | 0.377 | 0.377 | 0.037 | 1.000 | 0.511 | 0.512 | 0.038 | 1.000 |
| MR-Lasso | 0.229 | 0.229 | 0.020 | 1.000 | 0.302 | 0.301 | 0.032 | 1.000 | 0.458 | 0.456 | 0.056 | 1.000 |
| MR-Egger | 1.015 | 1.015 | 0.198 | 1.000 | 1.098 | 1.097 | 0.190 | 1.000 | 0.996 | 0.994 | 0.182 | 1.000 |
| Contamination Mixture | 0.207 | 0.207 | 0.020 | 1.000 | 0.224 | 0.223 | 0.030 | 1.000 | 0.340 | 0.284 | 0.164 | 1.000 |
| MR-Mix | 0.316 | 0.284 | 0.138 | 0.445 | 0.443 | 0.410 | 0.209 | 0.475 | 0.605 | 0.604 | 0.223 | 0.535 |
| MR-RAPS | 0.303 | 0.302 | 0.023 | 1.000 | 0.412 | 0.412 | 0.032 | 1.000 | 0.523 | 0.522 | 0.037 | 1.000 |

**Fig. 3:**
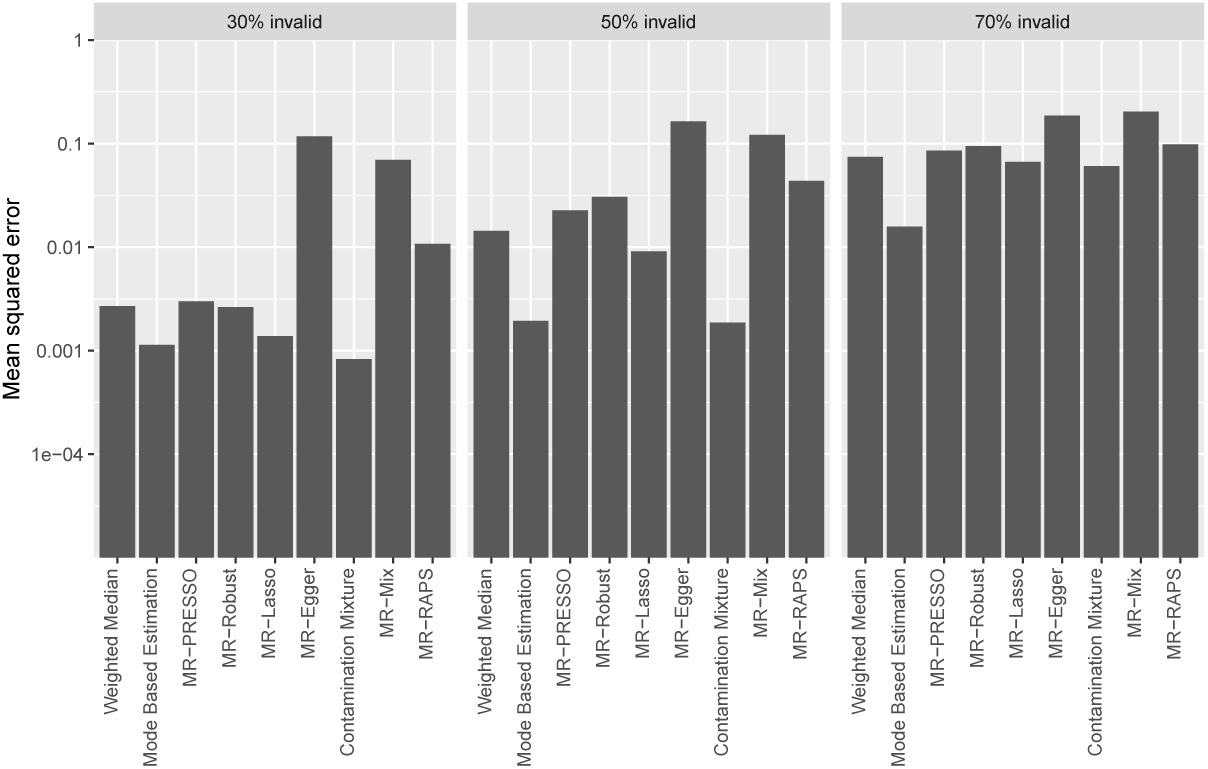
Mean squared errors for the different methods in scenario 2 (directional pleiotropy, InSIDE satisfied) with a null causal effect for 30 variants. Note the vertical axis is on a logarithmic scale.

**Fig. 4:**
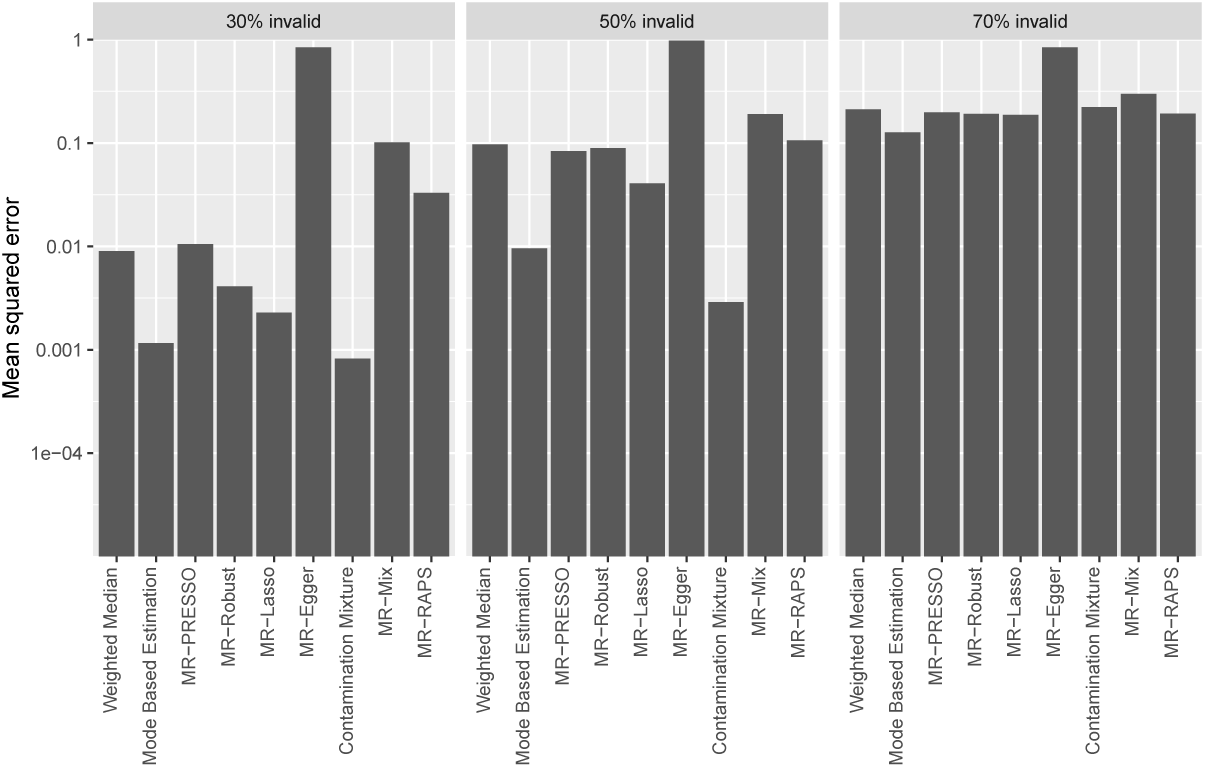
Mean squared errors for the different methods in scenario 3 (directional pleiotropy, InSIDE violated) with a null causal effect for 30 variants. Note the vertical axis is on a logarithmic scale.

**Fig. 5:**
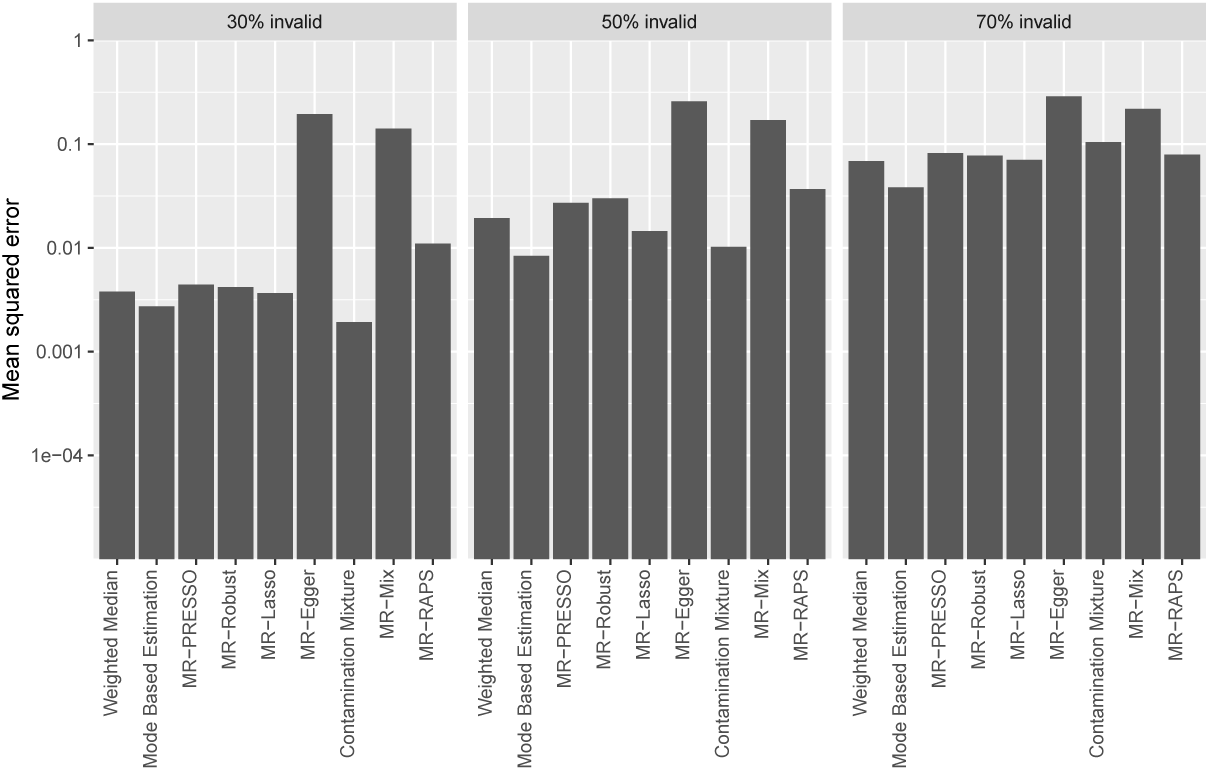
Mean squared error for the different methods in scenario 2 for 10 000 simulations, with directional pleiotropy and InSIDE satisfied with 10 variants.

**Fig. 6:**
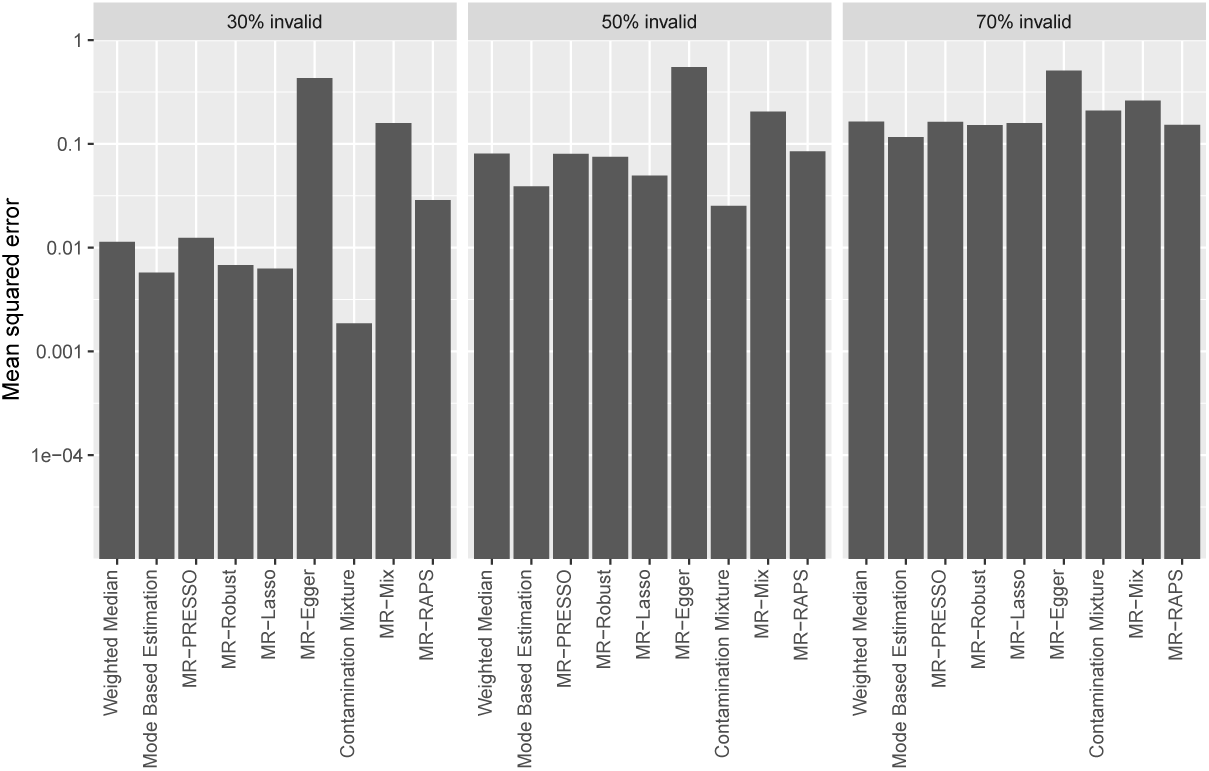
Mean squared error for the different methods in scenario 3 for 10 000 simulations, with directional pleiotropy and InSIDE violated with 10 variants.

**Fig. 7:**
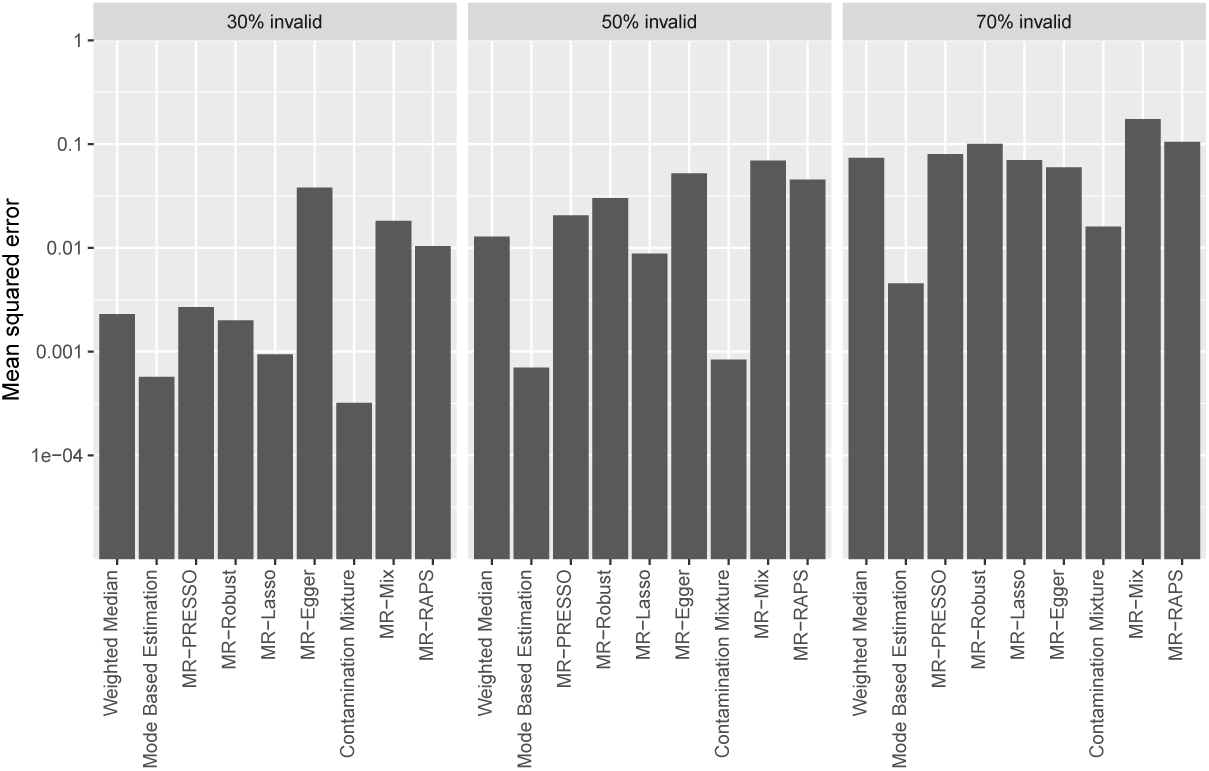
Mean squared error for the different methods in scenario 2 for 10 000 simulations, with directional pleiotropy and InSIDE satisfied with 100 variants.

**Fig. 8:**
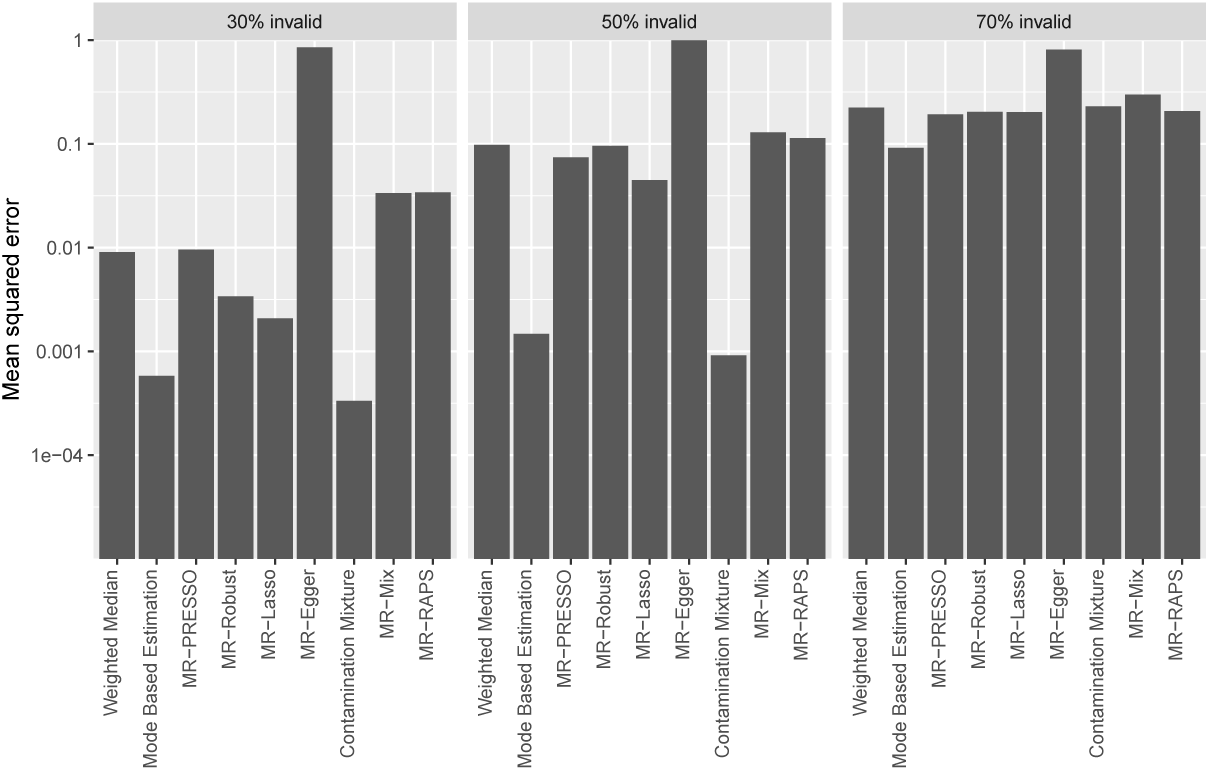
Mean squared error for the different methods in scenario 3 for 10 000 simulations, with directional pleiotropy and InSIDE violated with 100 variants.

Overall, judging by mean squared error, the contamination mixture and MBE methods performed best. The contamination mixture method performed slightly better with 30% and 50% invalid variants, and the MBE method performed better with 70% invalid variants. However, with some isolated exceptions, all the methods performed badly with 70% invalid instruments. Between these two methods, MBE tended to be more conservative, whereas the contamination mixture method had slightly lower standard deviation of estimates and increased power to detect a causal effect. Neither method consistently dominated the other in terms of Type 1 error rate. Several other methods performed well in particular scenarios.

Amongst consensus methods, estimates from the MBE method were less biased than those from the weighted median method, with lower Type 1 errors. The weighted median method had slightly higher power to detect a causal effect, although comparisons of power lose much of their value when a method has inflated Type 1 error rates. Amongst outlier-robust methods, estimates were similar amongst the methods, with the MR-Lasso method generally having the lowest bias, but MR-Robust having the lowest Type 1 error rates. None of the methods dominated in terms of power to detect a causal effect.

The modelling methods performed well in some scenarios, but less well in others. This is unsurprising, as in some scenarios, consistency assumptions for the methods were satisfied, and in others they were not. The MR-Egger method performed well in terms of Type 1 error rate in Scenarios 1 and 2, where the InSIDE assumption was satisfied. Estimates from the method were generally imprecise with low power. However, power in the MR-Egger method depends on the genetic associations with the exposure varying substantially between variants, which was not the case in the simulation study [38]. The contamination mixture method performed well with 30% and 50% valid instruments, with low bias and Type 1 error rates at or below 10% with 10 variants, 12% with 30 variants, and 20% with 100 variants. The MR-Mix method performed badly throughout, with highly inflated Type 1 error rates in almost all scenarios and comparatively low power to detect a causal effect. It performed slightly better with more genetic variants, although its performance was still worse than other methods. The MR-RAPS method performed well in Scenario 1, where its consistency assumption was satisfied, but less well in other scenarios with highly inflated Type 1 error rates.

### Empirical example: The effect of body mass index on coronary artery disease

Results from the empirical example are shown in Table 4. All methods agree that there is a positive effect of BMI on CAD risk, except for the MR-Mix method which gives a wide confidence interval that includes the null. The narrowest confidence intervals are for the outlier-robust methods (MR-Lasso, MR-Robust, MR-PRESSO), followed by the modelling methods except MR-Mix and MR-Egger (contamination mixture, MR-RAPS), then the consensus methods (weighted median, mode based estimation), and finally MR-Egger and MR-Mix.

**Table 4:**
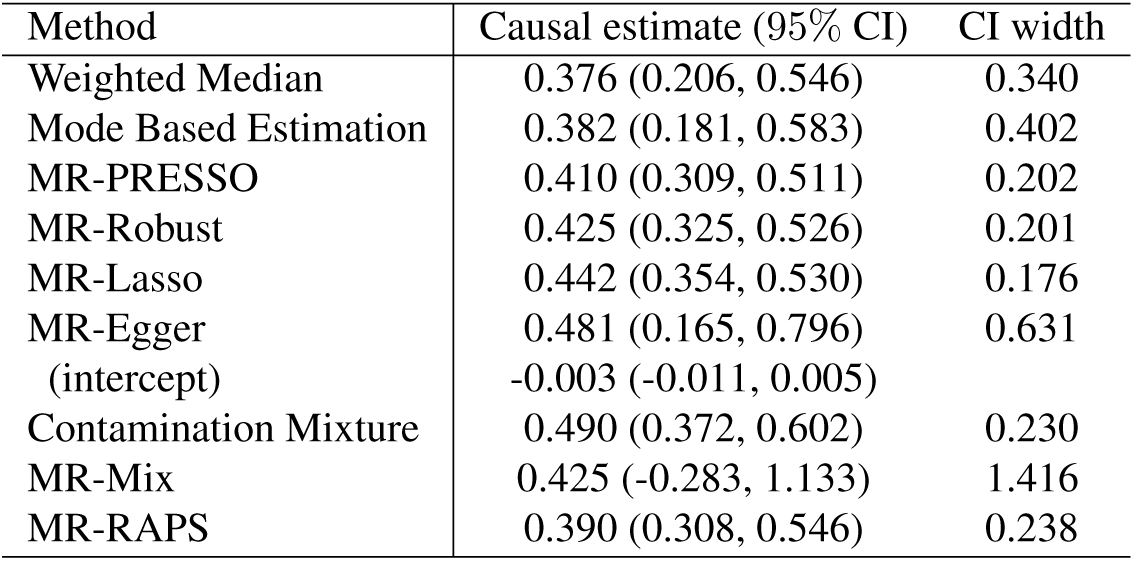
Estimates and 95% confidence intervals (CI) for the effect of BMI on coronary artery disease risk from robust methods. Estimates represent log odds ratios for CAD risk per 1 kg/m^2^ increase in BMI.

While the methods that detect outliers varied in terms of how lenient or strictly they identified outliers, they agreed on the order of outliers (Supplementary Table 8). The MR-Robust method was the most lenient, downweighting two variants as outliers. Each subsequent method in order of strictness identified all previously identified variants as outliers. MR-PRESSO excluded the two variants identified by MR-Robust plus an additional three variants. MR-RAPS identified these five plus an additional two variants. MR-Lasso identified an additional three variants, 10 in total. The contamination mixture method identified an additional 14 variants, 24 in total. MR-Mix identified an additional 21 variants, 45 in total. This suggests that any difference between results from outlier-robust methods are likely due to the strictness of outlier detection, rather than due to intrinsic differences in how the different methods select outliers. In several methods, the threshold at which outliers are detected can be varied by the analyst (for example, by varying the penalization parameter *λ* in MR-Lasso, or the significance threshold in MR-PRESSO). In practice, rather than performing different outlier-robust methods, it may be better to concentrate on one method, but vary this threshold.

## Discussion

In this paper, we have provided a review of robust methods for MR, focusing on methods that can be performed using summary data and implemented using standard statistical software. We have divided methods into three categories: consensus methods, outlier-robust methods, and modelling methods. Methods were compared in three ways: by their theoretical properties, including the assumptions required for the method to give a consistent estimate, in an extensive simulation study, and in an empirical investigation. A summary table comparing the methods is presented as Table 5.

**Table 5:**
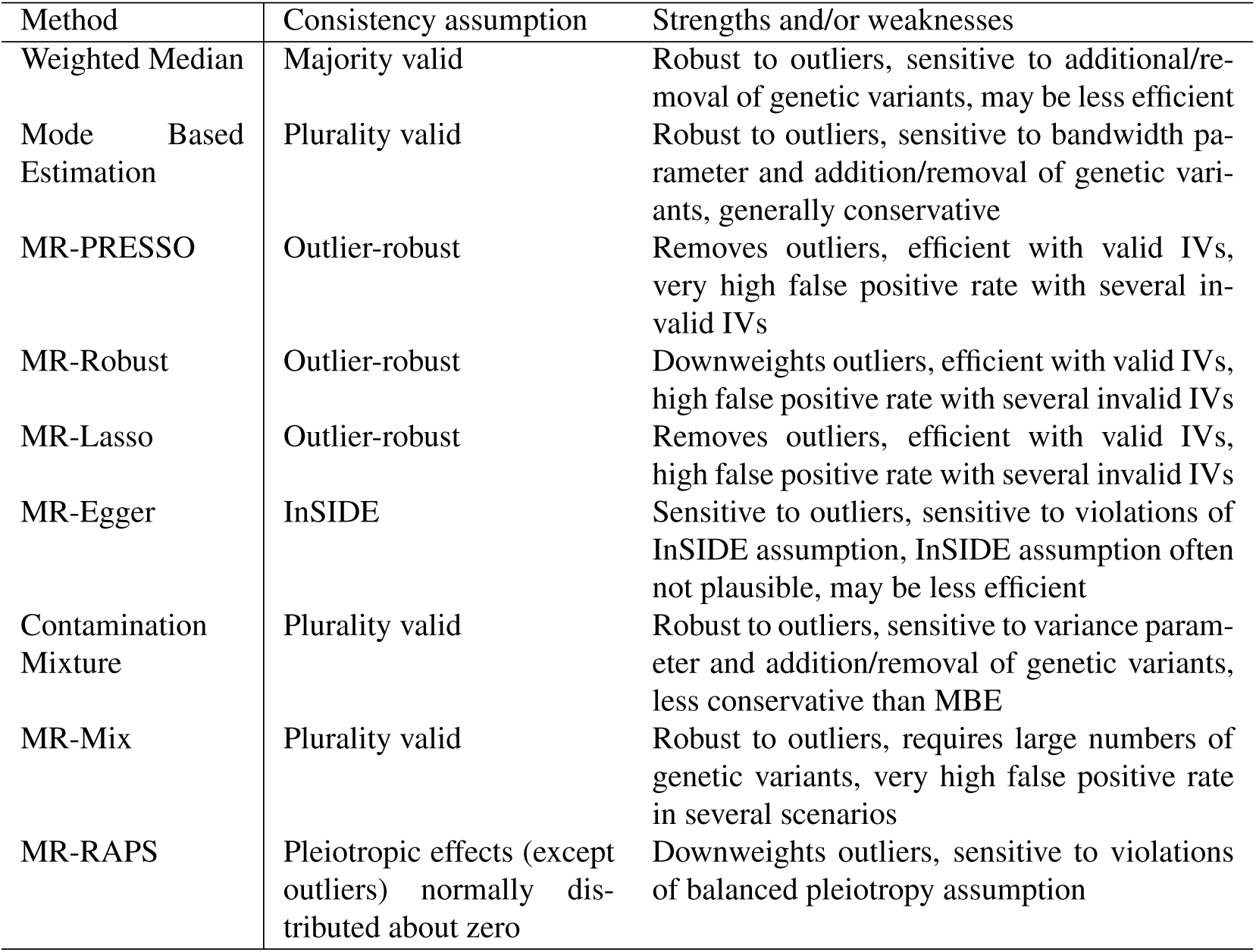
Summary comparison of methods.

While the use of robust methods for MR analyses with multiple genetic variants is highly recommended, it is not practical or desirable to perform and report results from every single robust method that has been proposed. Guidance is therefore needed as to which robust methods should be performed in practice. As an example, if an investigator performed the MR-PRESSO, MR-Robust, and MR-Lasso methods, they would have assessed robustness of the result to outliers, but they would not have not assessed other potential violations of the IV assumptions. The categorization of methods proposed here is not the only possible division of methods, but we hope it is practically useful. For instance, the contamination mixture and MR-Mix methods make the same ‘plurality valid’ assumption as the MBE method, and so could have been placed in the same category.

The similarity and ubiquity of the ‘outlier-robust’ and ‘majority/plurality valid’ assumptions should encourage investigators to consider methods that make alternative assumptions, such as the MR-Egger method. While the InSIDE assumption is often not plausible [38], the MR-Egger method and the intercept test have value in providing a different route to testing the validity of an MR study. Another potential choice is the constrained IV method, which uses information on measured confounders to construct a composite IV that is not associated with these confounders [12]. This method was not considered in the simulation study, as it requires additional data on confounders and individual participant data. Further methods development is needed to develop robust methods for summary data that make different consistency assumptions.

We encourage researchers to perform robust methods from different categories, and that make varied consistency assumptions. For example, an investigator could perform the weighted median method (majority valid assumption), the MBE and/or contamination mixture methods (both plurality valid assumption, MBE is more conservative), and the MR-Egger method (InSIDE assumption). If there are a few clear outliers in the data, then an outlier-robust method such as MR-PRESSO or MR-Robust could also be performed. While we are hesitant to make a definitive recommendation as each method has its own strengths and weaknesses, this set of methods would be a reasonable compromise between performing too few methods and not adequately assessing the IV assumptions, and performing so many methods that clarity is obscured. Another danger of the use of large numbers of methods is the possibility to cherry-pick results, either by an investigator seeking to present their results in a more positive light, or a reader picking the one method that gives a different result (such as the MR-Mix method in our empirical example).

One important limitation of these methods is the assumption that all valid IVs estimate the same causal effect. Particularly for complex risk factors such as BMI, it is possible that different genetic variants have different ratio estimates not because they are invalid IVs, but because there are different ways of intervening on BMI that lead to different effects on the outcome. This can be remedied somewhat in methods based on the IVW method by using a random-effects model [19], or in the contamination mixture method, where causal effects evidenced by different sets of variants will lead to a multimodal likelihood function, and potentially a confidence interval that consists of more than one region.

In summary, while robust methods for MR do not provide a perfect solution to violations of the IV assumptions, they are able to detect such violations and help investigators make more reliable causal inferences. Investigators should perform a range of robust methods that operate in different ways and make different assumptions to assess the robustness of findings from a MR investigation.

## Acknowledgements

We would like to thank Jack Bowden, Nilanjan Chatterjee, George Davey Smith, Ron Do, Christopher Foley, Fernando Hartwig, Gibran Hemani, Benjamin Neale, Nuala Sheehan, Dylan Small and Frank Windmeijer for input in selecting the scenarios and parameter values used in the simulation study. Eric Slob acknowledges funding from the Stichting Erasmus Trustfonds for his research visit to the MRC Biostatistics Unit. Stephen Burgess is supported by a Sir Henry Dale Fellowship jointly funded by the Wellcome Trust and the Royal Society (Grant Number 204623/Z/16/Z).

## Supporting Information

### S1 Details of simulation study

For each participant *i*, we simulate data on *J* genetic variants *G*_*i*1_, *G*_*i*2_, *…, G*_*iJ*_, a modifiable exposure *X*_*i*_, an outcome variable *Y*_*i*_, and a confounder *U*_*i*_ (assumed unknown). The confounder is a linear function of the genetic variants and an independent error term 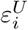. The effect of variant *j* on the confounder is represented by coefficient *φ*_*j*_. The exposure is linear in the genetic variants, the confounder and an independent error term 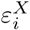. The effect of variant *j* on the exposure is represented by coefficient *γ*_*j*_ (this is zero for a valid IV). The outcome is linear in the genetic variants, exposure, confounders and an independent error term 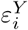. The effect of variant *j* on the outcome is represented by coefficient *α*_*j*_ (again, this is zero for a valid IV). The effect of the exposure on the outcome is represented by *θ*. The genetic variants are modelled as single nucleotide polymorphisms (SNPs) with minor allele frequency 30%, and take values 0, 1 or 2. The error terms 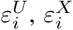 and 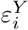 each follow an independent normal distribution with mean 0 and unit variance.

We can represent the model mathematically as:

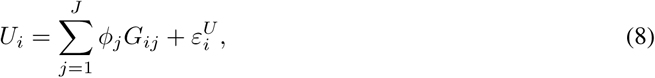

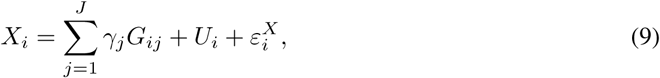

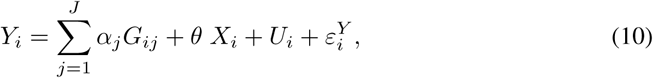

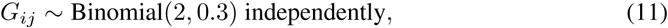

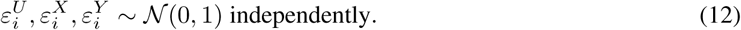

The causal effect of the exposure on the outcome was either taken as null (*θ* = 0) or positive (*θ* = 0.2). Genetic associations with the exposure *γ*_*j*_ are drawn from a truncated normal distribution. Parameters are chosen such that the total proportion of variance explained in the exposure by direct effects of the genetic variants is 10%. In scenario 3, the overall proportion of variance explained in the exposure by genetic variants is slightly larger, as there is an additional effect of the invalid IVs on the exposure via their effect on the confounder.

For valid IVs, *φ*_*j*_ = 0 and *α*_*j*_ = 0. For invalid IVs, in scenario 1 (balanced pleiotropy, InSIDE satisfied), the effects of the genetic variants on the outcome are generated from a normal distribution centered at zero (*α*_*j*_ ∼𝒩 (0, 0.15)) and genetic effects on the confounder are zero (*φ*_*j*_ = 0). In scenario 2 (directional pleiotropy, InSIDE satisfied), the effects of the genetic variants on the outcome are generated from a normal distribution centered away from zero (*α*_*j*_ ∼𝒩 (0.1, 0.075)) and genetic effects on the confounder are zero (*φ*_*j*_ = 0). In scenario 3 (directional pleiotropy, InSIDE violated), the direct effects of the genetic variants on the outcome are generated from a normal distribution centered away from zero (*α*_*j*_ ∼𝒩 (0.1, 0.075)) and genetic effects on the confounder are generated from a uniform distribution (*φ*_*j*_ ∼𝒰 (0, 0.1)).

Summary genetic association data are calculated by regression of the risk factor on each genetic variant in turn, and the outcome on each genetic variant in turn. Individual participant data are generated for 20 000 individuals: the exposure regressions are performed on the first 10 000 individuals, and the outcome regressions on the remaining 10 000 individuals. This represents a two-sample Mendelian randomization study. We generated 10 000 simulated datasets for each scenario, and for null and positive causal effects.

Each method is performed using the default options suggested by the authors of the method, either in the corresponding publication, or in the software code recommended by the authors. The weighted median method is performed using inverse-variance weights. The mode based estimation method is performed using inverse-variance weights, the ‘no measurement error’ assumption, and the default bandwidth setting (*φ* = 1). The MR-PRESSO method is performed using a significance cut-off of *p <* 0.05 for determining outliers. The MR-Lasso method is performed using the heterogeneity criterion for selecting the lasso penalty parameter. The contamination mixture method is performed using the standard deviation of the ratio estimates multiplied by 1.5 for the variance parameter. For MR-Mix, we choose an initial value of the probability mass at the null component as 0.6 and the initial value of the variance of the non-null component as 1 *×* 10^-5^. As the method performs a grid search, these decisions should not influence the results. For MR-RAPS, we use the overdispersed robust version with the Huber loss function. All regression models use random-effects.

The mean squared errors of the different methods are presented in Supplementary Figure 5 (10 variants, scenario 2), Supplementary Figure 6 (10 variants, scenario 3), Supplementary Figure 7 (100 variants, scenario 2), and Supplementary Figure 8 (100 variants, scenario 3). Note that in each case the vertical axis is on a logarithmic scale. Findings are similar to before among the different scenarios. We observe again that the performance of the mode based estimator is the best for the consensus based approach, MR-Robust gets the best result among the outlier-robust methods, and the contamination mixture approach has the best performance among the modelling methods.

In addition to the scenarios presented in the main paper, we also performed simulations with 500 genetic variants, and a wider range of wider range of proportions of invalid IVs (1%, 5%, and 10%). Due to computational burden, only 1000 simulated datasets were generated in each case.

The results of these additional simulations can be found in Table 6 (invalidness 1%, 5%, and 10%), and 7 (invalidness 30%, 50% and 70%).

**Table 6:** Mean, median, standard deviation (SD) of estimates, and empirical power (%) for scenarios with 500 genetic variants.

| Method | 1% invalid |  |  |  | 5% invalid |  |  |  | 10% invalid |  |  |  |
| --- | --- | --- | --- | --- | --- | --- | --- | --- | --- | --- | --- | --- |
|  | Mean | Median | SD | Power | Mean | Median | SD | Power | Mean | Median | SD | Power |
| Null causal effect: $\beta = 0$ | | | | | | | | | | | | |
| Scenario 1: Balanced pleiotropy, InSIDE satisfied |  |  |  |  |  |  |  |  |  |  |  |  |
| Weighted Median | 0.000 | 0.000 | 0.006 | 0.038 | 0.000 | 0.000 | 0.006 | 0.014 | 0.000 | 0.000 | 0.007 | 0.034 |
| Mode Based Estimation | 0.000 | -0.001 | 0.017 | 0.001 | 0.000 | 0.000 | 0.017 | 0.003 | 0.000 | 0.000 | 0.018 | 0.004 |
| MR-PRESSO | 0.000 | 0.000 | 0.005 | 0.049 | 0.000 | 0.000 | 0.005 | 0.046 | 0.000 | 0.000 | 0.006 | 0.059 |
| MR-Robust | 0.000 | 0.000 | 0.005 | 0.048 | 0.000 | 0.000 | 0.005 | 0.046 | 0.000 | 0.000 | 0.006 | 0.046 |
| MR-Lasso | 0.000 | 0.000 | 0.005 | 0.023 | 0.000 | 0.000 | 0.005 | 0.027 | 0.000 | 0.000 | 0.006 | 0.037 |
| MR-Egger | 0.000 | 0.000 | 0.023 | 0.049 | -0.001 | 0.000 | 0.036 | 0.047 | 0.000 | 0.000 | 0.047 | 0.054 |
| Contamination Mixture | 0.000 | 0.000 | 0.005 | 0.055 | 0.000 | 0.000 | 0.006 | 0.050 | 0.000 | 0.001 | 0.006 | 0.052 |
| MR-Mix | 0.000 | 0.000 | 0.023 | 0.007 | -0.001 | 0.000 | 0.020 | 0.004 | -0.001 | 0.000 | 0.021 | 0.005 |
| MR-RAPS | 0.000 | 0.000 | 0.005 | 0.047 | 0.000 | 0.000 | 0.006 | 0.016 | 0.000 | 0.000 | 0.007 | 0.020 |
| Scenario 2: Directional pleiotropy, InSIDE satisfied |  |  |  |  |  |  |  |  |  |  |  |  |
| Weighted Median | 0.001 | 0.001 | 0.006 | 0.021 | 0.005 | 0.005 | 0.006 | 0.073 | 0.011 | 0.011 | 0.007 | 0.289 |
| Mode Based Estimation | 0.000 | 0.000 | 0.016 | 0.003 | 0.000 | 0.000 | 0.017 | 0.004 | -0.001 | 0.000 | 0.037 | 0.000 |
| MR-PRESSO | 0.001 | 0.001 | 0.005 | 0.049 | 0.004 | 0.004 | 0.005 | 0.114 | 0.010 | 0.010 | 0.006 | 0.392 |
| MR-Robust | 0.000 | 0.000 | 0.005 | 0.041 | 0.002 | 0.002 | 0.005 | 0.072 | 0.006 | 0.006 | 0.006 | 0.152 |
| MR-Lasso | 0.001 | 0.001 | 0.005 | 0.046 | 0.003 | 0.003 | 0.005 | 0.078 | 0.005 | 0.006 | 0.006 | 0.184 |
| MR-Egger | 0.000 | 0.000 | 0.022 | 0.052 | 0.000 | 0.000 | 0.032 | 0.052 | 0.001 | 0.001 | 0.039 | 0.049 |
| Contamination Mixture | 0.000 | 0.000 | 0.005 | 0.048 | 0.001 | 0.001 | 0.006 | 0.061 | 0.002 | 0.002 | 0.006 | 0.067 |
| MR-Mix | 0.000 | 0.000 | 0.022 | 0.007 | 0.003 | 0.001 | 0.019 | 0.002 | 0.003 | 0.001 | 0.021 | 0.007 |
| MR-RAPS | 0.001 | 0.001 | 0.005 | 0.050 | 0.008 | 0.008 | 0.006 | 0.239 | 0.020 | 0.020 | 0.006 | 0.783 |
| Scenario 3: Directional pleiotropy, InSIDE violated |  |  |  |  |  |  |  |  |  |  |  |  |
| Weighted Median | 0.002 | 0.002 | 0.006 | 0.030 | 0.010 | 0.010 | 0.007 | 0.232 | 0.022 | 0.022 | 0.007 | 0.778 |
| Mode Based Estimation | 0.000 | 0.000 | 0.016 | 0.004 | 0.000 | 0.001 | 0.018 | 0.004 | 0.001 | 0.001 | 0.017 | 0.003 |
| MR-PRESSO | 0.001 | 0.000 | 0.005 | 0.059 | 0.004 | 0.004 | 0.005 | 0.130 | 0.013 | 0.013 | 0.007 | 0.582 |
| MR-Robust | 0.000 | 0.000 | 0.005 | 0.048 | 0.002 | 0.002 | 0.005 | 0.064 | 0.006 | 0.006 | 0.006 | 0.148 |
| MR-Lasso | 0.001 | 0.001 | 0.005 | 0.058 | 0.003 | 0.003 | 0.005 | 0.102 | 0.007 | 0.007 | 0.006 | 0.230 |
| MR-Egger | 0.031 | 0.030 | 0.027 | 0.269 | 0.140 | 0.139 | 0.043 | 0.956 | 0.250 | 0.249 | 0.051 | 0.999 |
| Contamination Mixture | 0.000 | 0.000 | 0.005 | 0.043 | 0.001 | 0.001 | 0.006 | 0.058 | 0.002 | 0.002 | 0.006 | 0.059 |
| MR-Mix | 0.002 | 0.000 | 0.018 | 0.007 | 0.004 | 0.001 | 0.018 | 0.003 | 0.005 | 0.000 | 0.018 | 0.002 |
| MR-RAPS | 0.002 | 0.002 | 0.005 | 0.063 | 0.014 | 0.014 | 0.006 | 0.487 | 0.035 | 0.035 | 0.006 | 0.998 |
| Positive causal effect: $\theta = +0.2$ | | | | | | | | | | | | |
| Scenario 1: Balanced pleiotropy, InSIDE satisfied |  |  |  |  |  |  |  |  |  |  |  |  |
| Weighted Median | 0.179 | 0.179 | 0.008 | 1.000 | 0.179 | 0.179 | 0.008 | 1.000 | 0.180 | 0.179 | 0.009 | 1.000 |
| Mode Based Estimation | 0.154 | 0.154 | 0.020 | 0.989 | 0.153 | 0.153 | 0.021 | 0.992 | 0.154 | 0.153 | 0.020 | 0.995 |
| MR-PRESSO | 0.188 | 0.188 | 0.007 | 1.000 | 0.188 | 0.188 | 0.007 | 1.000 | 0.189 | 0.189 | 0.008 | 1.000 |
| MR-Robust | 0.188 | 0.188 | 0.007 | 1.000 | 0.188 | 0.188 | 0.007 | 1.000 | 0.189 | 0.189 | 0.007 | 1.000 |
| MR-Lasso | 0.188 | 0.188 | 0.007 | 1.000 | 0.188 | 0.188 | 0.007 | 1.000 | 0.189 | 0.188 | 0.008 | 1.000 |
| MR-Egger | 0.010 | 0.009 | 0.027 | 0.073 | 0.009 | 0.009 | 0.038 | 0.057 | 0.010 | 0.009 | 0.049 | 0.055 |
| Contamination Mixture | 0.187 | 0.187 | 0.007 | 1.000 | 0.187 | 0.187 | 0.008 | 1.000 | 0.187 | 0.187 | 0.008 | 1.000 |
| MR-Mix | 0.060 | 0.055 | 0.034 | 0.076 | 0.057 | 0.052 | 0.033 | 0.012 | 0.048 | 0.045 | 0.033 | 0.002 |
| MR-RAPS | 0.200 | 0.200 | 0.007 | 1.000 | 0.199 | 0.199 | 0.008 | 1.000 | 0.200 | 0.199 | 0.008 | 1.000 |
| Scenario 2: Directional pleiotropy, InSIDE satisfied |  |  |  |  |  |  |  |  |  |  |  |  |
| Weighted Median | 0.181 | 0.181 | 0.008 | 1.000 | 0.186 | 0.186 | 0.009 | 1.000 | 0.193 | 0.193 | 0.009 | 1.000 |
| Mode Based Estimation | 0.155 | 0.154 | 0.020 | 0.993 | 0.155 | 0.155 | 0.022 | 0.992 | 0.156 | 0.156 | 0.021 | 0.991 |
| MR-PRESSO | 0.190 | 0.190 | 0.007 | 1.000 | 0.195 | 0.195 | 0.007 | 1.000 | 0.204 | 0.203 | 0.008 | 1.000 |
| MR-Robust | 0.189 | 0.189 | 0.007 | 1.000 | 0.192 | 0.192 | 0.007 | 1.000 | 0.198 | 0.197 | 0.008 | 1.000 |
| MR-Lasso | 0.189 | 0.189 | 0.007 | 1.000 | 0.192 | 0.192 | 0.007 | 1.000 | 0.196 | 0.196 | 0.008 | 1.000 |
| MR-Egger | 0.010 | 0.010 | 0.026 | 0.070 | 0.010 | 0.010 | 0.034 | 0.062 | 0.010 | 0.010 | 0.042 | 0.062 |
| Contamination Mixture | 0.187 | 0.187 | 0.008 | 1.000 | 0.188 | 0.189 | 0.008 | 1.000 | 0.189 | 0.190 | 0.009 | 1.000 |
| MR-Mix | 0.063 | 0.061 | 0.035 | 0.061 | 0.066 | 0.064 | 0.033 | 0.018 | 0.064 | 0.060 | 0.034 | 0.017 |
| MR-RAPS | 0.202 | 0.202 | 0.007 | 1.000 | 0.210 | 0.210 | 0.008 | 1.000 | 0.222 | 0.222 | 0.009 | 1.000 |
| Scenario 3: Directional pleiotropy, InSIDE violated |  |  |  |  |  |  |  |  |  |  |  |  |
| Weighted Median | 0.182 | 0.182 | 0.008 | 1.000 | 0.191 | 0.191 | 0.009 | 1.000 | 0.206 | 0.207 | 0.010 | 1.000 |
| Mode Based Estimation | 0.154 | 0.153 | 0.020 | 0.997 | 0.154 | 0.154 | 0.022 | 0.987 | 0.152 | 0.155 | 0.057 | 0.988 |
| MR-PRESSO | 0.190 | 0.189 | 0.007 | 1.000 | 0.195 | 0.195 | 0.008 | 1.000 | 0.211 | 0.210 | 0.009 | 1.000 |
| MR-Robust | 0.189 | 0.189 | 0.007 | 1.000 | 0.192 | 0.191 | 0.007 | 1.000 | 0.199 | 0.199 | 0.008 | 1.000 |
| MR-Lasso | 0.189 | 0.189 | 0.007 | 1.000 | 0.192 | 0.192 | 0.008 | 1.000 | 0.199 | 0.199 | 0.009 | 1.000 |
| MR-Egger | 0.044 | 0.043 | 0.031 | 0.360 | 0.161 | 0.160 | 0.046 | 0.965 | 0.278 | 0.277 | 0.054 | 1.000 |
| Contamination Mixture | 0.187 | 0.187 | 0.008 | 1.000 | 0.188 | 0.188 | 0.008 | 1.000 | 0.190 | 0.190 | 0.009 | 1.000 |
| MR-Mix | 0.060 | 0.058 | 0.033 | 0.057 | 0.053 | 0.049 | 0.031 | 0.016 | 0.043 | 0.040 | 0.028 | 0.008 |
| MR-RAPS | 0.203 | 0.202 | 0.007 | 1.000 | 0.215 | 0.215 | 0.008 | 1.000 | 0.240 | 0.240 | 0.009 | 1.000 |

**Table 7:** Mean, median, standard deviation (SD) of estimates, and empirical power (%) for scenarios with 500 genetic variants.

| Method | 30% invalid |  |  |  | 50% invalid |  |  |  | 70% invalid |  |  |  |
| --- | --- | --- | --- | --- | --- | --- | --- | --- | --- | --- | --- | --- |
|  | Mean | Median | SD | Power | Mean | Median | SD | Power | Mean | Median | SD | Power |
| Null causal effect: $\theta = 0$ | | | | | | | | | | | | |
| Scenario 1: Balanced pleiotropy, InSIDE satisfied |  |  |  |  |  |  |  |  |  |  |  |  |
| Weighted Median | 0.000 | -0.001 | 0.010 | 0.054 | -0.001 | 0.000 | 0.015 | 0.099 | 0.000 | -0.001 | 0.021 | 0.164 |
| Mode Based Estimation | 0.003 | -0.001 | 0.101 | 0.005 | -0.002 | -0.001 | 0.020 | 0.004 | -0.004 | -0.001 | 0.112 | 0.008 |
| MR-PRESSO | 0.000 | 0.000 | 0.010 | 0.080 | -0.001 | 0.000 | 0.016 | 0.150 | 0.000 | 0.000 | 0.023 | 0.187 |
| MR-Robust | 0.000 | -0.001 | 0.009 | 0.043 | 0.000 | 0.000 | 0.018 | 0.059 | -0.001 | -0.001 | 0.026 | 0.048 |
| MR-Lasso | 0.000 | 0.000 | 0.009 | 0.058 | -0.001 | -0.001 | 0.013 | 0.085 | 0.000 | -0.001 | 0.019 | 0.136 |
| MR-Egger | -0.001 | 0.000 | 0.076 | 0.049 | -0.001 | -0.001 | 0.097 | 0.053 | 0.001 | 0.000 | 0.115 | 0.052 |
| Contamination Mixture | 0.000 | 0.000 | 0.010 | 0.085 | -0.001 | -0.002 | 0.015 | 0.131 | 0.000 | 0.000 | 0.025 | 0.197 |
| MR-Mix | 0.001 | 0.000 | 0.016 | 0.002 | 0.000 | 0.000 | 0.020 | 0.000 | 0.000 | 0.000 | 0.015 | 0.000 |
| MR-RAPS | 0.000 | 0.000 | 0.013 | 0.018 | 0.000 | 0.001 | 0.021 | 0.040 | -0.001 | -0.001 | 0.028 | 0.050 |
| Scenario 2: Directional pleiotropy, InSIDE satisfied |  |  |  |  |  |  |  |  |  |  |  |  |
| Weighted Median | 0.105 | 0.105 | 0.014 | 1.000 | 0.294 | 0.292 | 0.027 | 1.000 | 0.440 | 0.440 | 0.025 | 1.000 |
| Mode Based Estimation | 0.006 | 0.006 | 0.016 | 0.002 | 0.035 | 0.035 | 0.021 | 0.052 | 0.207 | 0.183 | 0.100 | 0.762 |
| MR-Robust | 0.086 | 0.086 | 0.014 | 0.999 | 0.313 | 0.313 | 0.018 | 1.000 | 0.438 | 0.438 | 0.018 | 1.000 |
| MR-Lasso | 0.061 | 0.061 | 0.011 | 1.000 | 0.233 | 0.233 | 0.023 | 1.000 | 0.426 | 0.426 | 0.025 | 1.000 |
| MR-PRESSO | 0.121 | 0.120 | 0.014 | 1.000 | 0.285 | 0.284 | 0.019 | 1.000 | 0.424 | 0.424 | 0.020 | 1.000 |
| MR-Egger | 0.000 | 0.000 | 0.059 | 0.050 | 0.000 | 0.000 | 0.069 | 0.052 | 0.000 | 0.000 | 0.072 | 0.044 |
| Contamination Mixture | 0.011 | 0.011 | 0.010 | 0.254 | 0.042 | 0.041 | 0.020 | 0.797 | 0.502 | 0.527 | 0.120 | 1.000 |
| MR-Mix | 0.004 | 0.000 | 0.011 | 0.000 | 0.004 | 0.000 | 0.011 | 0.001 | 0.012 | 0.000 | 0.020 | 0.000 |
| MR-RAPS | 0.188 | 0.188 | 0.014 | 1.000 | 0.341 | 0.340 | 0.017 | 1.000 | 0.460 | 0.460 | 0.019 | 1.000 |
| Scenario 3: Directional pleiotropy, InSIDE violated |  |  |  |  |  |  |  |  |  |  |  |  |
| Weighted Median | 0.048 | 0.048 | 0.010 | 1.000 | 0.121 | 0.120 | 0.016 | 1.000 | 0.256 | 0.257 | 0.024 | 1.000 |
| Mode Based Estimation | 0.003 | 0.003 | 0.016 | 0.004 | 0.016 | 0.015 | 0.016 | 0.018 | 0.054 | 0.055 | 0.022 | 0.313 |
| MR-PRESSO | 0.063 | 0.062 | 0.010 | 1.000 | 0.161 | 0.160 | 0.015 | 1.000 | 0.284 | 0.285 | 0.019 | 1.000 |
| MR-Robust | 0.051 | 0.050 | 0.010 | 0.999 | 0.180 | 0.180 | 0.016 | 1.000 | 0.309 | 0.309 | 0.017 | 1.000 |
| MR-Lasso | 0.033 | 0.033 | 0.009 | 0.990 | 0.109 | 0.109 | 0.015 | 1.000 | 0.260 | 0.260 | 0.024 | 1.000 |
| MR-Egger | 0.485 | 0.485 | 0.061 | 1.000 | 0.537 | 0.537 | 0.061 | 1.000 | 0.473 | 0.473 | 0.062 | 1.000 |
| Contamination Mixture | 0.010 | 0.010 | 0.009 | 0.257 | 0.030 | 0.030 | 0.014 | 0.738 | 0.095 | 0.092 | 0.036 | 0.993 |
| MR-Mix | 0.003 | 0.000 | 0.020 | 0.003 | 0.006 | 0.000 | 0.026 | 0.003 | 0.013 | 0.000 | 0.038 | 0.004 |
| MR-RAPS | 0.103 | 0.103 | 0.011 | 1.000 | 0.216 | 0.215 | 0.015 | 1.000 | 0.328 | 0.328 | 0.017 | 1.000 |
| Positive causal effect: $\theta = +0.2$ | | | | | | | | | | | | |
| Scenario 1: Balanced pleiotropy, InSIDE satisfied |  |  |  |  |  |  |  |  |  |  |  |  |
| Weighted Median | 0.180 | 0.179 | 0.013 | 1.000 | 0.179 | 0.179 | 0.016 | 1.000 | 0.180 | 0.179 | 0.023 | 1.000 |
| Mode Based Estimation | 0.154 | 0.154 | 0.022 | 0.987 | 0.152 | 0.151 | 0.023 | 0.983 | 0.146 | 0.145 | 0.029 | 0.978 |
| MR-PRESSO | 0.189 | 0.189 | 0.012 | 1.000 | 0.188 | 0.188 | 0.018 | 1.000 | 0.190 | 0.189 | 0.025 | 1.000 |
| MR-Robust | 0.189 | 0.189 | 0.011 | 1.000 | 0.188 | 0.188 | 0.018 | 1.000 | 0.189 | 0.188 | 0.026 | 1.000 |
| MR-Lasso | 0.189 | 0.189 | 0.011 | 1.000 | 0.188 | 0.188 | 0.015 | 1.000 | 0.190 | 0.189 | 0.022 | 1.000 |
| MR-Egger | 0.010 | 0.010 | 0.078 | 0.056 | 0.011 | 0.009 | 0.098 | 0.056 | 0.011 | 0.012 | 0.115 | 0.051 |
| Contamination Mixture | 0.188 | 0.188 | 0.012 | 1.000 | 0.190 | 0.190 | 0.017 | 1.000 | 0.196 | 0.194 | 0.028 | 1.000 |
| MR-Mix | 0.021 | 0.000 | 0.028 | 0.001 | 0.006 | 0.000 | 0.019 | 0.000 | 0.002 | 0.000 | 0.015 | 0.000 |
| MR-RAPS | 0.200 | 0.200 | 0.014 | 1.000 | 0.200 | 0.200 | 0.022 | 1.000 | 0.201 | 0.201 | 0.028 | 1.000 |
| Scenario 2: Directional pleiotropy, InSIDE satisfied |  |  |  |  |  |  |  |  |  |  |  |  |
| Weighted Median | 0.307 | 0.307 | 0.018 | 1.000 | 0.488 | 0.487 | 0.026 | 1.000 | 0.619 | 0.618 | 0.026 | 1.000 |
| Mode Based Estimation | 0.164 | 0.163 | 0.022 | 0.986 | 0.201 | 0.202 | 0.105 | 0.971 | 0.395 | 0.384 | 0.097 | 0.983 |
| MR-PRESSO | 0.343 | 0.343 | 0.017 | 1.000 | 0.500 | 0.501 | 0.019 | 1.000 | 0.622 | 0.623 | 0.020 | 1.000 |
| MR-Robust | 0.314 | 0.314 | 0.018 | 1.000 | 0.510 | 0.510 | 0.018 | 1.000 | 0.627 | 0.627 | 0.019 | 1.000 |
| MR-Lasso | 0.273 | 0.273 | 0.015 | 1.000 | 0.449 | 0.448 | 0.024 | 1.000 | 0.613 | 0.614 | 0.025 | 1.000 |
| MR-Egger | 0.010 | 0.010 | 0.061 | 0.051 | 0.010 | 0.009 | 0.071 | 0.053 | 0.010 | 0.010 | 0.076 | 0.056 |
| Contamination Mixture | 0.207 | 0.207 | 0.015 | 1.000 | 0.288 | 0.280 | 0.051 | 1.000 | 0.717 | 0.721 | 0.081 | 1.000 |
| MR-Mix | 0.022 | 0.019 | 0.022 | 0.000 | 0.020 | 0.012 | 0.025 | 0.007 | 0.039 | 0.030 | 0.037 | 0.011 |
| MR-RAPS | 0.394 | 0.394 | 0.016 | 1.000 | 0.545 | 0.544 | 0.018 | 1.000 | 0.662 | 0.662 | 0.020 | 1.000 |
| Scenario 3: Directional pleiotropy, InSIDE violated |  |  |  |  |  |  |  |  |  |  |  |  |
| Weighted Median | 0.238 | 0.238 | 0.013 | 1.000 | 0.319 | 0.319 | 0.019 | 1.000 | 0.445 | 0.445 | 0.025 | 1.000 |
| Mode Based Estimation | 0.156 | 0.161 | 0.137 | 0.987 | 0.179 | 0.176 | 0.053 | 0.990 | 0.223 | 0.221 | 0.032 | 0.990 |
| MR-PRESSO | 0.273 | 0.273 | 0.013 | 1.000 | 0.375 | 0.374 | 0.018 | 1.000 | 0.490 | 0.489 | 0.020 | 1.000 |
| MR-Robust | 0.259 | 0.259 | 0.013 | 1.000 | 0.382 | 0.382 | 0.018 | 1.000 | 0.500 | 0.500 | 0.019 | 1.000 |
| MR-Lasso | 0.234 | 0.234 | 0.012 | 1.000 | 0.321 | 0.321 | 0.018 | 1.000 | 0.461 | 0.461 | 0.025 | 1.000 |
| MR-Egger | 0.535 | 0.535 | 0.064 | 1.000 | 0.592 | 0.591 | 0.065 | 1.000 | 0.527 | 0.526 | 0.066 | 1.000 |
| Contamination Mixture | 0.204 | 0.204 | 0.012 | 1.000 | 0.238 | 0.237 | 0.020 | 1.000 | 0.386 | 0.368 | 0.087 | 1.000 |
| MR-Mix | 0.039 | 0.036 | 0.034 | 0.009 | 0.034 | 0.016 | 0.048 | 0.011 | 0.052 | 0.027 | 0.069 | 0.036 |
| MR-RAPS | 0.308 | 0.308 | 0.013 | 1.000 | 0.419 | 0.418 | 0.018 | 1.000 | 0.530 | 0.531 | 0.020 | 1.000 |

With few invalid IVs, most methods had reasonable behaviour. An exception was the MR-Egger method, which had inflated Type 1 error rates in Scenario 3 even with only 1% of variants invalid. Although we would expect the outlier-removal methods to behave best with few invalid IVs, in fact most methods have some mechanism for providing robustness to outliers, and so it was difficult to differentiate between the methods.

### S2 Outliers according to different methods

**Table 8:** Genetic variants identified as outliers by the different methods in the Mendelian Randomization study of the effect of BMI on cardiovascular disease risk.

| Variant | MR-Robust | MR-PRESSO | MR-RAPS | MR-Lasso | Contamination mixture | MR-Mix |
| --- | --- | --- | --- | --- | --- | --- |
| rs11191560 | ✓ | ✓ | ✓ | ✓ | ✓ | ✓ |
| rs2075650 | ✓ | ✓ | ✓ | ✓ | ✓ | ✓ |
| rs2176040 |  | ✓ | ✓ | ✓ | ✓ | ✓ |
| rs6567160 |  | ✓ | ✓ | ✓ | ✓ | ✓ |
| rs7903146 |  | ✓ | ✓ | ✓ | ✓ | ✓ |
| rs11727676 |  |  | ✓ | ✓ | ✓ | ✓ |
| rs17024393 |  |  | ✓ | ✓ | ✓ | ✓ |
| rs11126666 |  |  |  | ✓ | ✓ | ✓ |
| rs13078960 |  |  |  | ✓ | ✓ | ✓ |
| rs9914578 |  |  |  | ✓ | ✓ | ✓ |
| rs1000940 |  |  |  |  | ✓ | ✓ |
| rs11057405 |  |  |  |  | ✓ | ✓ |
| rs11847697 |  |  |  |  | ✓ | ✓ |
| rs12446632 |  |  |  |  | ✓ | ✓ |
| rs12566985 |  |  |  |  | ✓ | ✓ |
| rs16907751 |  |  |  |  | ✓ | ✓ |
| rs205262 |  |  |  |  | ✓ | ✓ |
| rs2650492 |  |  |  |  | ✓ | ✓ |
| rs2836754 |  |  |  |  | ✓ | ✓ |
| rs3849570 |  |  |  |  | ✓ | ✓ |
| rs4787491 |  |  |  |  | ✓ | ✓ |
| rs492400 |  |  |  |  | ✓ | ✓ |
| rs7243357 |  |  |  |  | ✓ | ✓ |
| rs9641123 |  |  |  |  | ✓ | ✓ |
| rs10938397 |  |  |  |  |  | ✓ |
| rs10968576 |  |  |  |  |  | ✓ |
| rs11030104 |  |  |  |  |  | ✓ |
| rs11688816 |  |  |  |  |  | ✓ |
| rs12016871 |  |  |  |  |  | ✓ |
| rs13021737 |  |  |  |  |  | ✓ |
| rs13191362 |  |  |  |  |  | ✓ |
| rs13201877 |  |  |  |  |  | ✓ |
| rs1460676 |  |  |  |  |  | ✓ |
| rs1516725 |  |  |  |  |  | ✓ |
| rs1528435 |  |  |  |  |  | ✓ |
| rs17203016 |  |  |  |  |  | ✓ |
| rs2176598 |  |  |  |  |  | ✓ |
| rs2287019 |  |  |  |  |  | ✓ |
| rs2820292 |  |  |  |  |  | ✓ |
| rs3810291 |  |  |  |  |  | ✓ |
| rs3817334 |  |  |  |  |  | ✓ |
| rs543874 |  |  |  |  |  | ✓ |
| rs7164727 |  |  |  |  |  | ✓ |
| rs7599312 |  |  |  |  |  | ✓ |
| rs7899106 |  |  |  |  |  | ✓ |

### S3 Software code

This section includes the code to run the robust methods used in this paper. Please note that the MR-Mix package is not publicly available, please contact the authors for the package ^1^.

~~~
*#install required packages*
**if** (**!require**(“MendelianRandomization”)) {**install.packages**(“
 MendelianRandomization”)} **else** {}
**if** (**!require**(“mr.raps”)) {**install.packages**(“mr.raps”)} **else** {}
**if** (**!require**(“devtools”)) {**install.packages**(“devtools”)} **else** {}
**if** (**!require**(“penalized”)) {**install.packages**(“penalized”)} **else** {}

**library**(“devtools”)
devtools::**install_**github(“rondolab**/**MR-PRESSO”)

*#load packages*
**library**(“MendelianRandomization”)
**library**(“mr.raps”)
**library**(“MRMix”)
**library**(“MRPRESSO”)
**library**(“penalized”)

*#create dataframe and object by different methods*
mr**_frame<-as.data.frame**(**cbind**(ldlc,ldlcse,chdlodds,chdloddsse))
**names**(mr**_frame**)**<-c**(“ldlc”,“ldlcse”,“chdlodds”,“chdloddsse”)
mr**_**object**<-**mr**_**input(bx = ldlc, bxse = ldlcse, **by** = chdlodds, byse =
   chdloddsse)*#create used by methods from MendelianRandomization package*

*#perform weighted median*
mr**_median**(mr**_**object,weighting = “weighted”, iterations = 10000)

*#perform Mode based estimation*
mr**_**mbe(mr**_**object, weighting = “weighted”, stderror = “delta”, phi = 1,
   seed = 19940407, iterations = 10000, distribution = “normal”, alpha = 0.05)
*#perform MR-PRESSO*
mr**_**presso(BetaOutcome = “chdlodds”, BetaExposure = “ldlc”,
SdOutcome = “chdloddsse”, SdExposure = “ldlcse”, OUTLIERtest = TRUE, DISTORTIONtest = TRUE, **data** = mr**_frame**, NbDistribution = 1000, SignifThreshold = 0.05)

*#perform MR-Robust*
mr**_**ivw(mr**_**object,“random”, robust = TRUE)

*#define function for MR-Lasso with heterogeneity criterion*
MR**_**lasso**<-function**(betaYG,betaXG,sebetaYG){

  betaYGw = betaYG**/**sebetaYG *# dividing the association estimates by sebetaYG is equivalent*
  betaXGw = betaXG**/**sebetaYG *# to weighting by sebetaYG^-2*
  pleio = **diag**(**rep**(1, **length**(betaXG)))
  l1grid = **c**(**seq**(from=0.1, to=5, **by**=0.1), **seq**(from=5.2, to=10, **by** =0.2))
  *# values of lambda for grid search*
  l1grid**_**rse = NULL; l1grid**_length** = NULL; l1grid**_beta** = NULL;
    l1grid**_se** = NULL
  **for** (i in 1:**length**(l1grid)) {
  l1grid**_which** = **which**(**attributes**(penalized(betaYGw, pleio,
    betaXGw, lambda1=l1grid[i], **trace**=FALSE))**$**penalized==0)
  l1grid**_**rse[i] = **summary**(**lm**(betaYG[l1grid**_which**]**∼**betaXG[l1grid**_**
    **which**]-1, **weights**=sebetaYG[l1grid**_which**]^-2))**$**sigma
  l1grid**_length**[i] = **length**(l1grid**_which**)
  l1grid**_beta**[i] = **lm**(betaYG[l1grid**_which**]**∼**betaXG[l1grid**_which**]-1,
    **weights**=sebetaYG[l1grid**_which**]^-2)**$coef**[1]
  l1grid**_se**[i] = **summary**(**lm**(betaYG[l1grid**_which**]**∼**betaXG[l1grid**_**
  **which**]-1, **weights**=sebetaYG[l1grid**_which**]^-2))**$coef**[1,2]**/min**(
 **summary**(**lm**(betaYG[l1grid**_which**]**∼**betaXG[l1grid**_which**]-1,
 **weights**=sebetaYG[l1grid**_which**]^-2))**$**sigma, 1)
}
l1which**_**hetero = **c**(**which**(l1grid**_**rse[1:(**length**(l1grid)-1)]>1**& diff**(
 l1grid**_**rse)>**qchisq**(0.95, **df**=1)**/**l1grid**_length**[2:**length**(l1grid)])
, **length**(l1grid))[1]
*# heterogeneity criterion for choosing lambda*

l1hetero**_beta** = l1grid**_beta**[l1which**_**hetero]
l1hetero**_se** = l1grid**_se**[l1which**_**hetero]
**list**(ThetaEstimate=l1hetero**_beta**, ThetaSE=l1hetero**_se)**

}

*#perform MR-Lasso*
MR**_**lasso(mr**_frame$**chdlodds,mr**_frame$**ldlc,mr**_frame$**chdloddsse)

*#perform MR-Egger*
mr**_**egger(mr**_**object)

*#define function for contamination mixture*
contaminationmixture**<-function**(**by**,bx,byse){
 iters = 2001; theta = **seq**(from=-3, to=3, **by**=2**/**(iters-1))
 *# if the causal estimate (and confidence interval) is not expected*
    *to lie between −1 and 1 then change from and to (and maybe increase iters)*
 ratio = **by/**bx; ratio.**se** = **abs**(byse**/**bx); psi = 1.5**\*sd**(ratio)
 lik=NULL
 **for** (j1 in 1:iters) {
   lik.inc = **exp**(−(theta[j1]-ratio)^2**/**2**/**ratio.**se**^2) **/sqrt**(2**\***pi**\***
      ratio.**se**^2)
   lik.exc = **exp**(−ratio^2**/**2**/**(psi^2+ratio.**se**^2)) **/**(**sqrt**(2**\***pi**\***(psi^2+
      ratio.**se**^2)))
   valid = (lik.inc>lik.exc)**\***1
   lik[j1] = **prod**(**c**(lik.inc[valid==1], lik.exc[valid==0]))
   **if** (**which.max**(lik)==**length**(lik)) {valid.best = valid}
}
 phi = **ifelse**(**sum**(valid.best)<1.5, 1, **max**(**sqrt**(**sum**(((ratio[valid. best==1]-**weighted.mean**(ratio[valid.best==1], ratio.**se**[valid. best==1]^-2))^2 **\*** ratio.**se**[valid.best==1]^-2)) **/**(**sum**(valid.best)-1)), 1))
 loglik = **log**(lik)
 whichin = **which**(2**\***loglik>(2**\*max**(loglik)-**qchisq**(0.95, **df**=1)**\***phi^2))
 theta[**which.max**(loglik)] *# estimate*
 theta[whichin[1]] *# lower limit of CI*
 theta[whichin[**length**(whichin)]] *# upper limit of CI*
**list**(ThetaEstimate=theta[**which.max**(loglik)], ThetaLower=theta[whichin[1]], ThetaUpper= theta[whichin[**length**(whichin)]])
}

*#perform contamination mixture, note we removed the 27th variable due to having a ratio to close to infty.*
contaminationmixture(mr**_frame$**chdlodds[−27],mr**_frame$**ldlc[−27],mr**_ frame$**chdloddsse[−27])

*#perform MR-Mix*
estMix = MRMix(mr**_frame$**chdlodds, mr**_frame$**ldlc, mr**_frame$** chdloddsse^2, mr**_frame$**ldlcse^2)
**se** = MRMix**_se**(mr**_frame$**chdlodds, mr**_frame$**ldlc, mr**_frame$**chdloddsse
 ^2, mr**_frame$**ldlcse^2, estMix**$**theta, estMix**$**pi0, estMix**$**sigma2)

*#perform MR-RAPS with Huber loss function*

mr.raps.overdispersed.robust(mr**_frame$**chdlodds, mr**_frame$**ldlc, mr**_**
**frame$**chdloddsse, mr**_frame$**ldlcse,
           loss.**function** = “huber”, k = 1.345,
               initialization = **c**(“l2”), suppress.**warning**
                 = FALSE, diagnosis = FALSE, niter = 20,
                tol = .**Machine$double**.eps^0.5)
~~~

## Footnotes

1 current maintainer of the package is Guanghao Qi.

